# Ecological and evolutionary drivers of hemoplasma infection and bacterial genotype sharing in a Neotropical bat community

**DOI:** 10.1101/2019.12.21.885921

**Authors:** Daniel J. Becker, Kelly A. Speer, Alexis M. Brown, M. Brock Fenton, Alex D. Washburne, Sonia Altizer, Daniel G. Streicker, Raina K. Plowright, Vladimir E. Chizhikov, Nancy B. Simmons, Dmitriy V. Volokhov

## Abstract

Most emerging pathogens can infect multiple species, underscoring the importance of understanding the ecological and evolutionary factors that allow some hosts to harbor greater infection prevalence and share pathogens with other species. However, our understanding of pathogen jumps is primarily based around viruses, despite bacteria accounting for the greatest proportion of zoonoses. Because bacterial pathogens in bats (Order: Chiroptera) can have conservation and human health consequences, studies that examine the ecological and evolutionary drivers of bacterial prevalence and barriers to pathogen sharing are crucially needed. We here studied hemotropic *Mycoplasma* spp. (i.e., hemoplasmas) across a species-rich bat community in Belize over two years. Across 469 bats spanning 33 species, half of individuals and two-thirds of species were hemoplasma positive. Infection prevalence was higher for males and for species with larger body mass and colony sizes. Hemoplasmas displayed high genetic diversity (21 novel genotypes) and strong host specificity. Evolutionary patterns supported co-divergence of bats and bacterial genotypes alongside phylogenetically constrained host shifts. Bat species centrality to the network of shared hemoplasma genotypes was phylogenetically clustered and unrelated to prevalence, further suggesting rare—but detectable—bacterial sharing between species. Our study highlights the importance of using fine phylogenetic scales when assessing host specificity and suggests phylogenetic similarity may play a key role in host shifts for not only viruses but also bacteria. Such work more broadly contributes to increasing efforts to understand cross-species transmission and epidemiological consequences of bacterial pathogens.

## Introduction

Most pathogens that cause disease in humans, domestic animals, and wildlife are capable of infecting multiple host species (Woolhouse, Taylor, & Haydon, 2001). However, predicting which hosts maintain pathogens and identifying their role in cross-species transmission can be challenging, as many hosts can be infected but not play key roles in the reservoir community (A. Fenton, Streicker, Petchey, & Pedersen, 2015). Pathogen jumps between species depends on infection prevalence in the donor host, transmission opportunities between donor and recipient species, and suitability of the recipient host for pathogen replication (Plowright et al., 2017). Each of these steps can be shaped by ecological and evolutionary factors (VanderWaal & Ezenwa, 2016). For example, small-bodied species can have greater competence, the ability to transmit new infections, than larger species (Downs, Schoenle, Han, Harrison, & Martin, 2019), and host switching is often constrained by phylogeny, owing to similarity in immunological barriers to pathogen replication between closely related species (Streicker et al., 2010). Identifying the ecological and evolutionary factors that allow some species to harbor greater prevalence and have facilitated pathogen sharing can improve our general understanding of disease emergence (Fountain-Jones et al., 2018). Examining evolutionary associations between hosts and pathogens can further uncover factors favoring host shifts versus codivergence and assess the frequency of cross-species transmission (Geoghegan, Duchêne, & Holmes, 2017).

Given the public health and agricultural burdens of many zoonotic pathogens such as SARS coronavirus, avian influenza virus, and rabies virus, many investigations of pathogen prevalence and emergence focus on viruses (Geoghegan et al., 2017; Olival et al., 2017). However, more zoonoses are caused by bacteria than other pathogen taxa (Han, Kramer, & Drake, 2016), and bacterial pathogens can negatively impact newly infected host species (e.g., *Mycoplasma galliscepticum*, a poultry pathogen, caused rapid population declines in wild house finches; Hochachka & Dhondt, 2000). More attention to bacteria and their propensity for host specificity versus generalism is accordingly important for understanding whether factors that govern cross-species transmission of viruses can be extended to other pathogens (Bonneaud, Weinert, & Kuijper, 2019). Bacterial pathogens have been especially understudied for bats (Mühldorfer, 2013), in contrast to intensive studies of zoonotic viruses across the Chiroptera (Luis et al., 2015). However, many bacterial pathogens are likely important to both bat conservation and human health due to pathogenic effects on bats themselves as well as their zoonotic potential (Becker et al., 2018; Evans, Bown, Timofte, Simpson, & Birtles, 2009).

To determine the ecological and evolutionary drivers of bacterial prevalence and barriers to pathogen sharing, we focused on hemotropic *Mycoplasma* spp. (i.e., hemoplasmas) in a species-rich bat community in Belize (Fenton et al., 2001; Herrera, Duncan, Clare, Fenton, & Simmons, 2018). The Neotropics have remarkable bat diversity owing to adaptive radiation in the Phyllostomidae (Gunnell & Simmons, 2012), producing a range of feeding strategies (e.g., frugivory, carnivory, sanguivory), body sizes, and roost preferences (Monteiro & Nogueira, 2011). Hemoplasmas are intracellular erythrocytic bacteria transmitted through direct contact (Cohen et al., 2018; Museux et al., 2009) and also possibly arthropod vectors (Willi, Boretti, Meli, et al., 2007). Hemoplasmas can cause acute and chronic anemia, especially for immunocompromised hosts; however, many animals develop inapparent infections and are asymptomatic (Messick, 2004). As *Mycoplasma* spp. lack many of the metabolic pathways associated with energy production and synthesis of cell components found in other bacteria, they are fully dependent on host cells (Citti & Blanchard, 2013). Hemoplasmas have therefore been described as mostly host specialists (Pitcher & Nicholas, 2005), yet interspecies and potentially zoonotic transmission can occur (Maggi et al., 2013; Willi, Boretti, Tasker, et al., 2007). Hemoplasmas are common and genetically diverse in bats (Di Cataldo, Kamani, Cevidanes, Msheliza, & Millán, 2020; Ikeda et al., 2017; Mascarelli et al., 2014; Millán et al., 2019, 2019; Volokhov et al., 2017), which offers an ideal model system for identifying the ecological and evolutionary factors structuring bacterial infection risks within and between host species.

Many cross-species comparisons of pathogen infection risks and sharing have used less-diverse host communities (Johnson et al., 2012; VanderWaal, Atwill, Isbell, & McCowan, 2014) or global datasets of host–pathogen associations that can be limited by heterogeneous sampling effort and variation in pathogen detection methods (Dallas et al., 2019; Huang, Bininda-Emonds, Stephens, Gittleman, & Altizer, 2014). Our focus on a widespread pathogens group in a highly diverse host community allowed us to capitalize on strong host trait variation while controlling for sampling effort and diagnostics methods (Becker, Crowley, Washburne, & Plowright, 2019; Han, Kramer, et al., 2016). Past work has also used host–pathogen networks to characterize contemporary or historic transmission at often coarse taxonomic scales (e.g., pathogen species complexes or genera; Blyton, Banks, Peakall, Lindenmayer, & Gordon, 2014; VanderWaal et al., 2014). However, as bat species can be infected by multiple hemoplasma genotypes, and because genotypes with ≥ 99% sequence identity of their 16S rRNA genes can represent different bacterial species (Volokhov, Hwang, Chizhikov, Danaceau, & Gottdenker, 2017; Volokhov, Simonyan, Davidson, & Chizhikov, 2012), focusing on genotypes provides finer-scale resolution to determine the ecological and evolutionary features of species that facilitate pathogen sharing and to identify likely maintenance hosts of bacterial infections (Fountain-Jones et al., 2018).

We asked three questions about the relative contribution of ecological traits and evolutionary history to structuring infection patterns and pathogen sharing. First, what are the individual and ecological predictors of hemoplasma infection in a Neotropical bat community? Second, does the distribution of hemoplasma genotypes across the bat community map onto the bat phylogeny, as predicted by host–pathogen codivergence? Lastly, if genotype sharing among host species occurs, which host clades or traits best predict species ability to share pathogens? We predicted that ecological covariates such as ectoparasitism and large host colonies could increase bacteria risk through vector-borne or density-dependent transmission (McCallum, Barlow, & Hone, 2001; Willi, Boretti, Meli, et al., 2007). We also expected hemoplasma genotypes would be specific to particular host species but that more closely related bats would share hemoplasma genotypes, indicating phylogenetically restricted host shifts (Pitcher & Nicholas, 2005). However, ecological traits that increase the risk of pathogen exposure between species, such as occupying a greater diversity of roosting habitats, could also facilitate pathogen genotype sharing among less closely related hosts (McKee et al., 2019).

## Methods

### Bat capture and sampling

From April 24 to May 6 2017 and from April 23 to May 5 2018, we sampled 469 bats from 33 species captured in two adjacent areas in the Orange Walk District of Belize: Lamanai Archaeological Reserve (LAR) and Ka’Kabish (KK; Fig. 1). The LAR is bordered by the New River Lagoon, forest, and agriculture, while KK is a remnant patch of forest surrounded by agriculture located 10 km away. We consider these sites to be independent, as the small home ranges of many Neotropical bat species and lack of continuous forest between LAR and KK has likely restricted most individual bat inter-site movement (Jones, Hämsch, Page, Kalko, & O’mara, 2017; Loayza & Loiselle, 2008); however, some species may have greater connectivity (e.g., inter-roost movement of *Desmodus rotundus* occurs rarely). At least 44 of the 70 bat species in Belize have been recorded in this region (Herrera et al., 2018; Reid, 1997). Bats were captured with mist nets primarily along flight paths and occasionally at the exits of roosts from 19:00 until 22:00. Harp traps were also set from 18:00 to 05:00. In KK, *Desmodus rotundus*, *Trachops cirrhosus*, and *Chrotopterus auritus* were captured at their shared roost. In the LAR, *Desmodus rotundus*, *Saccopteryx bilineata*, and *Glossophaga soricina* were sampled from a shared roost, although individuals were also sampled along flight paths across the greater site. More broadly, patterns of roost sharing of bat species in northern Belize remain elusive.

**Figure 1.**
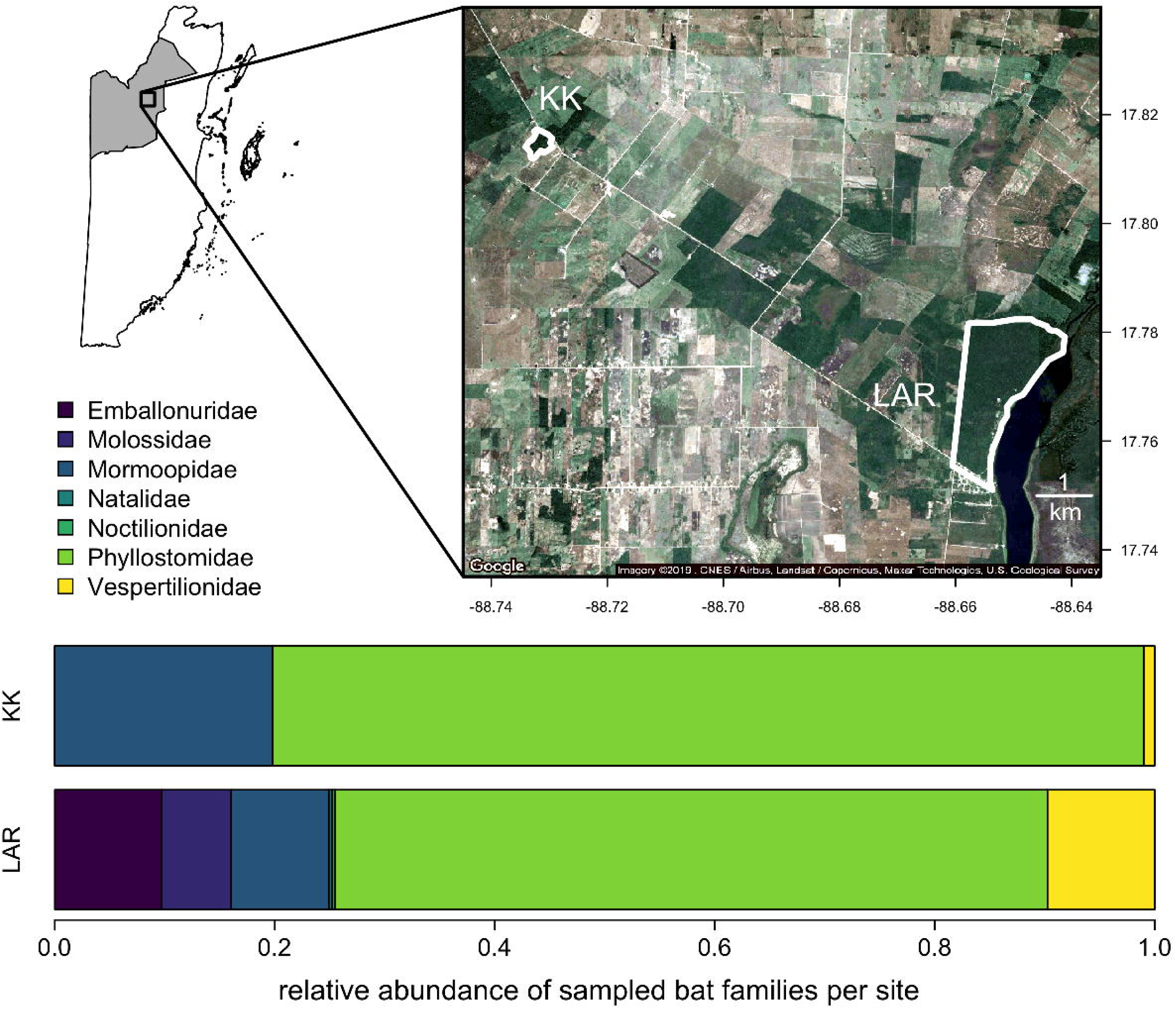
Study sites in northern Belize. The shaded inset shows the location of Orange Walk District. Borders show the boundaries of the LAR (Lamanai Archaeological Reserve) and KK (Ka’Kabish). White and tan shading indicates agricultural and urban development, while dark green shading represents intact forest. Satellite imagery was derived from Google Maps. Stacked bar plots display the relative abundance of each sampled bat family per study site.

Bats were placed in cloth bags until processing and were identified to species (and sex) based on morphology (Reid, 1997). Reproductive activity was indicated by the presence of scrotal testes in males and by the evidence of pregnancy or lactation in females; across bat species, 69% of males and 42% of females were in reproductive condition. We also visually screened bats for the presence of ectoparasites (i.e., bat flies, ticks, bat bugs, mites; Ter Hofstede, Fenton, & Whitaker, 2004). We collected 3–30 μL of blood (volumes were dependent on bat mass) by lancing the propatagial vein with a sterile needle. Blood was collected with heparinized capillary tubes and stored on Whatman FTA cards to preserve bacterial DNA. Field procedures followed guidelines for safe and humane handling of bats published by of the American Society of Mammalogists (Sikes, Care, & Mammalogists, 2016) and were approved by the Institutional Animal Care and Use Committees of the University of Georgia (A2014 04-016-Y3-A5) and American Museum of Natural History (AMNHIACUC-20170403 and AMNHIACUC-20180123). Fieldwork was authorized by the Belize Forest Department under permits WL/2/1/17(16), WL/2/1/17(19), and WL/2/1/18(16). Sample size was similar between years (2017=202, 2018=267) but varied by site (LAR=365, KK=101). More species were sampled for blood at LAR (*n*=33) than KK (*n*=17; Fig. 1), reflecting site differences in species richness (Herrera et al., 2018). We sampled 1–139 individuals per bat species (the maximum being *Desmodus rotundus*), with a mean of 14 individuals per bat species (Table S1).

### DNA extraction, PCR amplification, and amplicon sequencing

Genomic DNA was extracted from blood on FTA cards using QIAamp DNA Investigator Kits (Qiagen). We tested DNA for hemoplasmas using PCR with primers and procedures described in prior analyses (Volokhov et al., 2017). We included blank FTA punches as an extraction control, ultrapure water as a negative control, and *Candidatus* Mycoplasma haemozalophi DNA as a positive control (Volokhov et al., 2011). Amplicons from PCR-positive samples were purified by electrophoresis and extracted using the QIAquick Gel Extraction Kit (Qiagen).

To determine hemoplasma infection status, all 16S rRNA amplicons were directly sequenced by Macrogen (https://www.macrogenusa.com/). Amplicons were sequenced with the same primers used for PCR amplification and then with internal (walking) primers when needed (Volokhov et al., 2017). Negative DNA samples were tested for amplification quality using the universal PCR primers targeting the mammal mitochondrial 16S rRNA gene (Volokhov, Kong, George, Anderson, & Chizhikov, 2008) or the mitochondrial cytochrome *c* oxidase subunit 1 gene (Clare, Lim, Engstrom, Eger, & Hebert, 2007); all hemoplasma-negative DNA samples gave positive signal in the mitochondrial 16S rRNA gene-and/or the COI-specific PCR assays. All amplified sequences were subjected to chimeric sequence analysis using DECIPHER (Wright, Yilmaz, & Noguera, 2012) and UCHIME (Edgar, Haas, Clemente, Quince, & Knight, 2011). All hemoplasma sequences have been deposited in GenBank under accession numbers MH245119–MH245194 and MK353807–MK353892; four positive samples were identified as *Bartonella* spp. during sequencing and were considered hemoplasma negative in our analyses.

### Bat phylogenetic data

We used the *rotl* and *ape* packages in R to extract a bat phylogeny from the Open Tree of Life and to calculate branch lengths with Grafen’s method (Michonneau, Brown, & Winter, 2016; Paradis, Claude, & Strimmer, 2004). To assess hemoplasma genotype sharing as a function of host phylogenetic similarity, we derived pairwise phylogenetic distances between the 33 sampled bat species (Fig. S1). Because more evolutionarily distant and distinct species could display less frequent bacterial genotype sharing owing to ecological and immunological barriers to pathogen exposure and replication (Huang, Drake, Gittleman, & Altizer, 2015), we used the *picante* package and our bat phylogeny to derive evolutionary distinctiveness (Kembel et al., 2010).

### Host species trait data

We obtained species-level data on host traits relevant to pathogen transmission from previously published sources (Table S2). We obtained fecundity (litter size, litters per year), body mass, and diet from the Amniote Life History and EltonTraits databases (Myhrvold et al., 2015; Wilman et al., 2014). For foraging ecology, which could affect bacterial exposure (e.g., trophic interactions; Kellner et al., 2018), we defined three dietary guilds: frugivory (including nectarivory; *n*=11), insectivory (*n*=18), and carnivory (including sanguivory and piscivory, *n*=4; González-Salazar, Martínez-Meyer, & López-Santiago, 2014). We also considered the proportion of plant-based items in diet. We simplified foraging strata into aerial (*n*=14), arboreal (*n*=16, including scansorial), and ground-or aquatic-level foraging (*n*=3). We also expanded prior compilations of wing aspect ratios and roost preferences to serve as proxies for ecological overlap among species (Fenton et al., 2001; Herrera et al., 2018; Reid, 1997). Roost type was simplified to open (e.g., only foliage; *n*=6) or closed (e.g., hollows, caves; *n*=27), and roost flexibility was simplified to using one (*n*=16) or multiple roost types (*n*=17). We classified maximum colony sizes as small-to-medium (e.g., under 100 individuals; *n*=20) or large (e.g., hundreds to thousands; *n*=13; Reid, 1997; Santana, Dial, Eiting, & Alfaro, 2011), as most values were reported in ranges. We did not record pairwise sympatry (e.g., Luis et al., 2015; McKee et al., 2019) given that all species occur in Belize (Fig. S2). Yet because more widely distributed species could have more opportunities for pathogen sharing due to range overlap, we used the *geosphere* package and data from the International Union for Conservation of Nature to derive geographic range size (Baillie, Hilton-Taylor, & Stuart, 2004; Hijmans, Williams, Vennes, & Hijmans, 2019). Missing species-level traits were taken from other databases, primary literature, or closely related species (Table S2).

### Individual-level analysis of bat infection status

We first used the *prevalence* package to estimate hemoplasma infection prevalence and its 95% confidence interval (Wilson method). We then used phylogenetic generalized linear mixed models (GLMMs) to test if infection status varied by sex, reproductive status, year, site, and ectoparasite presence while accounting for bat phylogenetic relatedness. We fit candidate GLMMs that considered all fixed effects as well as interactions between sex and reproduction and between site and year. We also considered a model that excluded site to account for possible non-independence of LAR and KK alongside an intercept-only model. As vampire bats were banded for a mark–recapture study (Volokhov et al., 2017) and some were sampled between and within years (*n*=14), we randomly selected one of each recapture. After removing recaptures and missing values (*n*=323), we fit the phylogenetic GLMMs using the *brms* package, default priors, and infection status as a Bernoulli-distributed response. We included random effects for bat species and phylogeny, the latter of which used the phylogenetic covariance matrix (Bürkner, 2017). We ran four chains for 20,000 iterations with a burn-in period of 10,000, thinned every 10 steps, for a total 4,000 samples. We compared GLMMs using the leave-one-out cross-validation (LOOIC) and assessed fit with a Bayesian *R^2^,* including the total modeled variance and that attributed to only the fixed effects (Gelman, Goodrich, Gabry, & Vehtari, 2019; Vehtari, Gelman, & Gabry, 2017). We then estimated fixed effects (means and 95% highest density intervals [HDI]) from the posterior distributions of each predictor from the top GLMM. Lastly, we assessed the sensitivity of individual-level analyses to overrepresentation of *Desmodus rotundus* by randomly subsampling this species by the maximum *n* for other bat species.

### Species-level analysis of hemoplasma prevalence

We next calculated infection prevalence per species, using the *metafor* package to estimate logit-transformed proportions and sampling variances (Viechtbauer, 2010). We used the *nlme* package to estimate phylogenetic signal as Pagel’s λ with a weighted phylogenetic generalized least squares (PGLS) model to account for within-species variance (Garamszegi, 2014). We next used a graph-partitioning algorithm, phylogenetic factorization, to flexibly identify clades with significantly different prevalence estimates at various taxonomic depths. We used the *taxize* package to obtain a bat taxonomy from the National Center for Biotechnology Information (Chamberlain & Szöcs, 2013) and used the *phylofactor* package to partition prevalence as a Bernoulli-distributed response in a GLM (Washburne et al., 2019). We determined the number of significant bat clades using Holm’s sequentially rejective test with a 5% family-wise error rate.

To identify species trait correlates of prevalence, we fit 11 PGLS models (weighted by sampling variance) with body mass, annual fecundity (litters per year * pups per litter), dietary guild, quantitative diet, foraging strata, aspect ratio, roost type, roost flexibility, colony size, geographic range size, and evolutionary distinctiveness as predictors. We also fit PGLS models with only sample size or an intercept. We compared models with Akaike information criterion corrected for small sample sizes (AICc) and estimated *R^2^* (Burnham & Anderson, 2002).

### Hemoplasma phylogenetic analyses and genotype assignment

We compared our 16S rRNA sequences to those in GenBank (Volokhov et al., 2017; Volokhov et al., 2011). Briefly, we aligned sequences using Clustal X, and inter-and intra-species similarity values were generated using BioEdit. Genetic distances were calculated with the Kimura two-parameter and Tamura–Nei models, and the phylogeny was constructed using MEGA X with the minimum evolution algorithm (Kumar, Stecher, Li, Knyaz, & Tamura, 2018).

We assigned hemoplasma genotypes to positive bats based on analysis of the 16S rRNA partial gene (860–1000 bp) sequences in GenBank and their clustering on the phylogeny. Genotypes were designated as novel if (*i*) sequences differed from the closest hemoplasma sequences in GenBank by ≥ 1.5% and/or (*ii*) if sequence similarity was <1.5% but genotype-specific reproducible mutations (at least two per sequence) were observed between hemoplasma sequences from at least two independent bat samples and the nearest GenBank hemoplasma sequences. These genotype-specific mutations were further used to differentiate closely related hemoplasma genotypes from our sample. We caution that genotype is not synonymous with species, as analysis of the 16S rRNA gene alone is insufficient for accurate species identification of *Mycoplasma* spp. (Volokhov et al., 2012). Future studies using genomics or housekeeping genes may identify independent but closely related hemoplasma species in our genotypes.

To assess if hemoplasma genotype assignments were associated with site and year, we used χ^2^ tests with *p* values generated through a Monte Carlo procedure. Prior to our phylogenetic and network analyses of genotype distributions across bat species (see below), we used another χ^2^ test to assess the association between hemoplasma genotype identify and bat host identity.

### Evolutionary relationships between bats and hemoplasmas

To determine the degree to which bat hemoplasma genotypes display host specificity and to describe their evolutionary relationships with host species, we used our bat and hemoplasma phylogenies to construct a binary association matrix. To test the dependence of the hemoplasma phylogeny upon the bat phylogeny and thus assess evidence of evolutionary codivergence, we applied the Procrustes Approach to Cophylogeny (PACo) using distance matrices and the *paco* package (Hutchinson, Cagua, Balbuena, Stouffer, & Poisot, 2017). We used a jackknife procedure to estimate the degree to which each bat–genotype link supported a hypothesis of phylogenetic congruence; links were supported if their upper 95% confidence interval was below the mean of all squared jackknife residuals (Balbuena, Míguez-Lozano, & Blasco-Costa, 2013).

### Hemoplasma genotype sharing among bat species

We used hemoplasma genotype assignments to create a network, with each node representing a bat species and edges representing shared genotypes among bat species pairs. We built an adjacency matrix using the *igraph* package and used the Louvain method to assess the structure of bat–hemoplasma communities within this network (Csardi & Nepusz, 2006). To test whether the distribution of hemoplasma genotypes across our Neotropical bat species is shaped by host phylogeny, we used two GLMs to predict counts of shared genotypes (Poisson errors) and the presence of sharing (binomial errors) by phylogenetic distance between bat species. We assessed statistical significance with a quadratic assignment procedure via the *sna* package (Butts, 2008).

We calculated two metrics of network centrality to quantify different aspects of how important a node (bat species) is to hemoplasma genotype sharing: degree and eigenvector centrality (Bell, Atkinson, & Carlson, 1999). Whereas degree indicates the number of other species with which a host shares bacterial genotypes (i.e., links per node), eigenvector centrality indicates the tendency for a host to share genotypes with species that also share more genotypes (i.e., connectivity). Eigenvector centrality is thus an extension of degree that can identify hubs of parasite sharing (Gómez, Nunn, & Verdú, 2013). These two metrics were moderately correlated (ρ=0.59), with many non-zero degree species displaying zero eigenvector centrality. To examine spatial and temporal patterns in host centrality, we built separate adjacency networks per each site and year. We fit separate GLMs to ask how hemoplasma sharing centrality was predicted by site, year, and the two-way interaction. Degree was modeled as a Poisson-distributed response, while eigenvector centrality was logit-transformed and used Gaussian errors. We next applied phylogenetic factorization to both metrics and weighted the algorithms by the square-root sample size per species (Garamszegi, 2014). We then fit the same PGLS models used in our prevalence analysis to identify the most competitive trait predictors of bat species centrality to hemoplasma sharing. To lastly assess whether network centrality is associated with hemoplasma prevalence, we fit two weighted PGLS models with each centrality metric as a univariate predictor.

## Results

### Hemoplasma infection status

We detected sequence-confirmed hemoplasma infection in 239 of 469 individuals (51%; 95% CI: 46–55%), with positive individuals in 23 of the 33 sampled bat species (Table S3). The most parsimonious phylogenetic GLMM explained 22% of the modeled variance in infection status and included only sex, reproductive status, ectoparasites, and year (ΔLOOIC=0.16, *w_i_*=0.27; Table S4), suggesting site to be uninformative. Males had higher odds of infection than females (OR=2.35, 95% HDI: 1.36–4.03), but risk was unrelated to ectoparasitism, reproduction, or year (Fig. 2A). This result was weakly sensitive to overrepresentation by *Desmodus rotundus*, as effect size and the prevalence sex difference was weaker when we randomly subsampled this species (Fig. 2). Across all models, fixed effects only explained up to 7% of the modeled variance, suggesting more variation explained by the species and phylogeny random effects.

**Figure 2.**
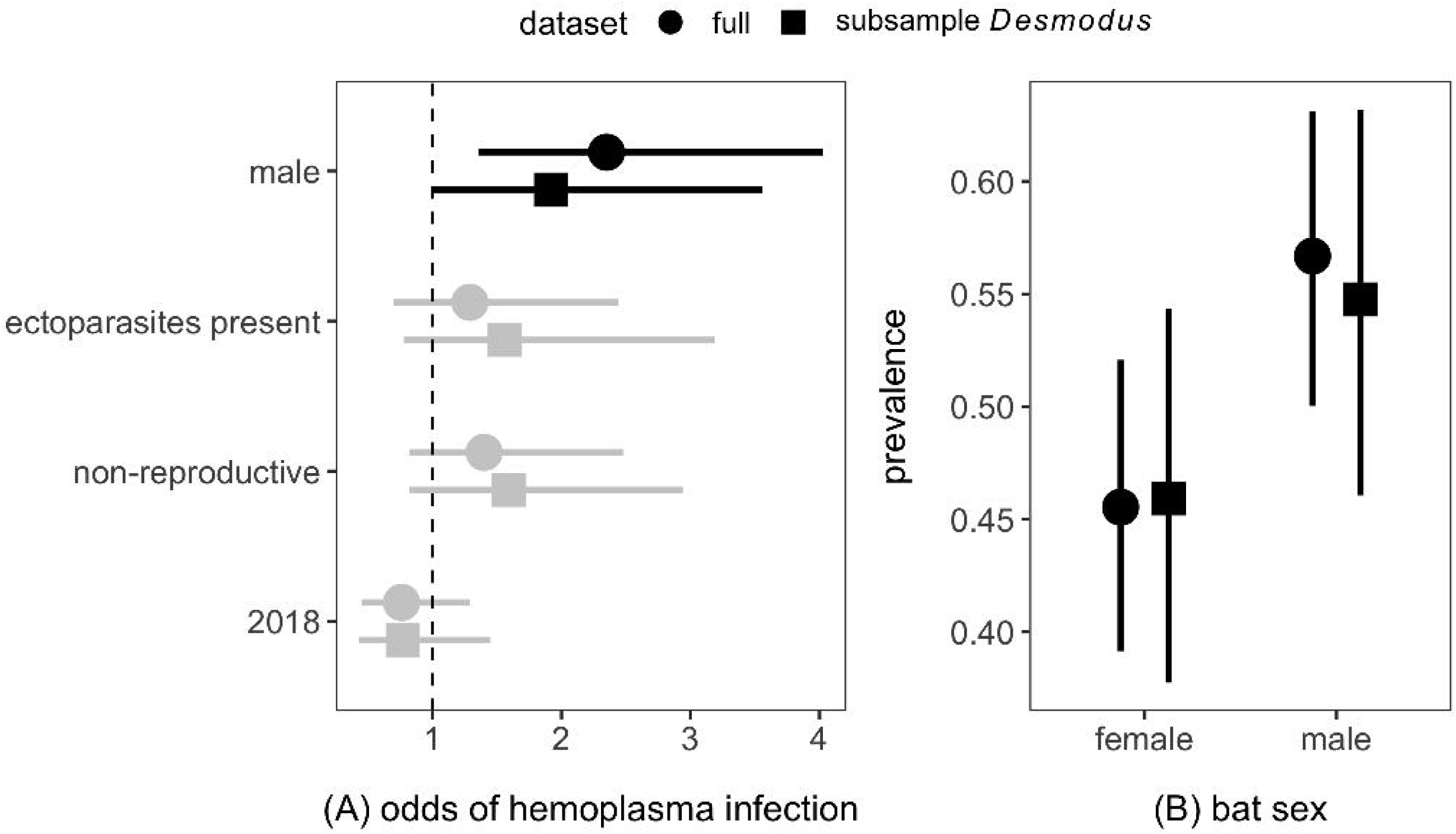
Predictors of individual bat hemoplasma infection status. (A) Odds ratios and 95% HDIs from the most parsimonious phylogenetic GLMM (Table S4). Estimates that do not overlap with 1 (dashed line) are displayed in black. Reference levels for the odds ratios include bats sampled at LAR, females, reproductive bats, absence of ectoparasites, and bats sampled in 2017. (B) Infection prevalence and 95% confidence intervals (Wilson method) stratified by sex. Results are shown for the full dataset and after randomly subsampling *Desmodus rotundus*.

### Inter-species variation in hemoplasma prevalence

Across bat species, hemoplasma prevalence ranged from 0% to 100% (x =0.37). We estimated Pagel’s λ in logit-transformed prevalence to be 0.39, indicating moderate phylogenetic signal. Similarly, phylogenetic factorization identified one bat clade with significantly lower prevalence compared to the paraphyletic remainder: the Emballonuridae (12% infected; Fig. 3A). Our trait-based analysis showed that relatively larger species (*β* =1.48, *p*=0.01, *R^2^*=0.24) and those with larger colonies (*β*=0.67, *p*=0.06, *R^22^*=0.20) had higher prevalence (Fig. 3B; Table 1). Relatively heavier (≥20 g) and larger colony species included *Desmodus rotundus*, *Molossus rufus*, and *Pteronotus mesoamericanus*, for which prevalence was greater than 58%. Although these three species were also heavily sampled, other well-sampled species such as *Sturnira parvidens* and *Carollia sowelli* had lower prevalence, and sample size did not predict prevalence (Table 1).

**Figure 3.**
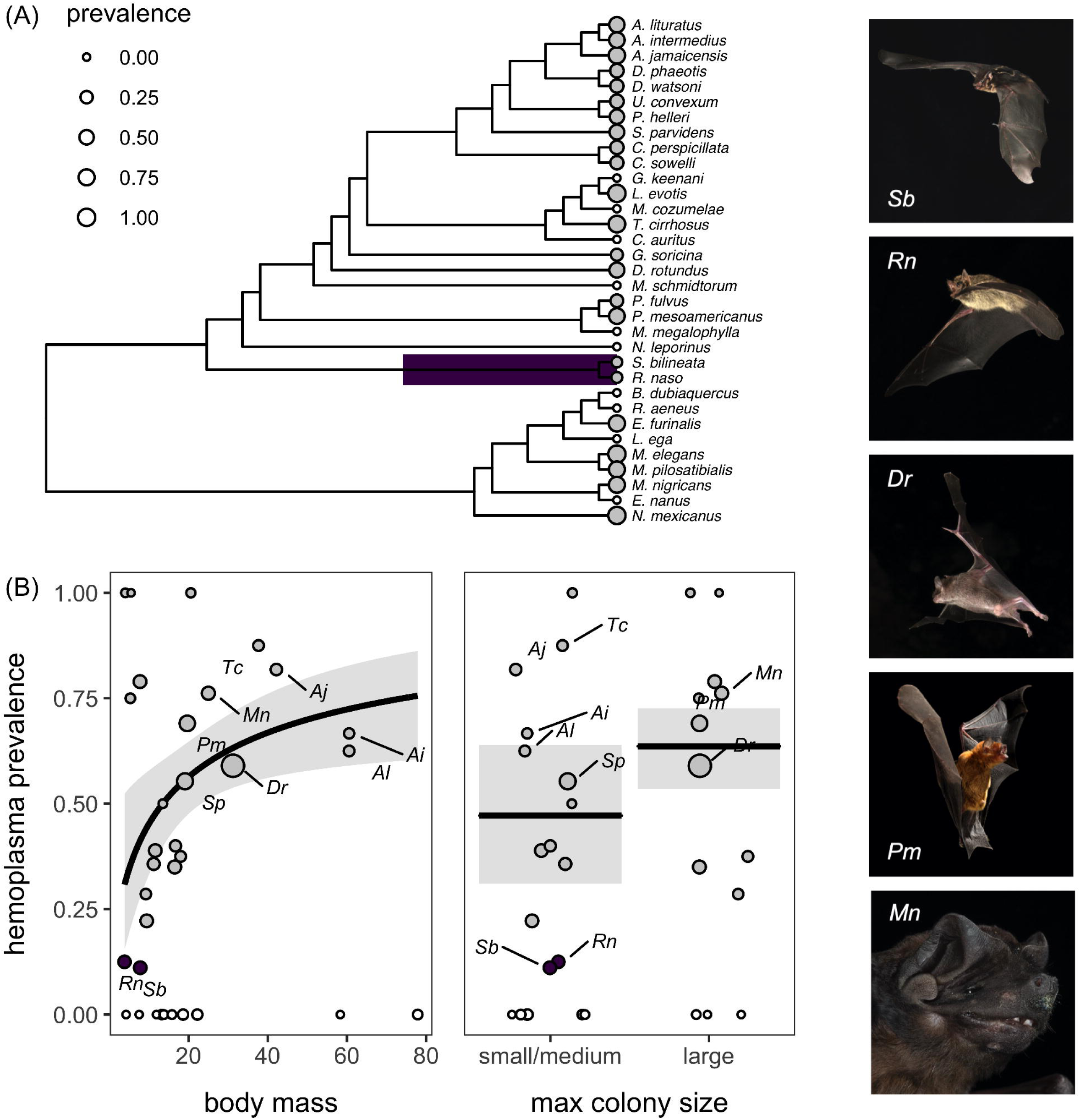
Predictors of species-level hemoplasma prevalence across the Belize bat community. (A) Clades with significantly different prevalence are highlighted. (B) Results from the top PGLS models predicting prevalence as a function of mass and colony size. Model fit and 95% confidence intervals are shown overlaid with data scaled by sample size; species from the clade identified through phylogenetic factorization are colored as in panel A. Species identified by phylogenetic factorization (*Saccopteryx bilineata* and *Rhynchonycteris naso*) and with larger body mass, colony size, and hemoplasma prevalence (*Desmodus rotundus*, *Molossus nigricans*, and *Pteronotus mesoamericanus*) are shown to the right (photographs by Brock Fenton).

**Table 1.**
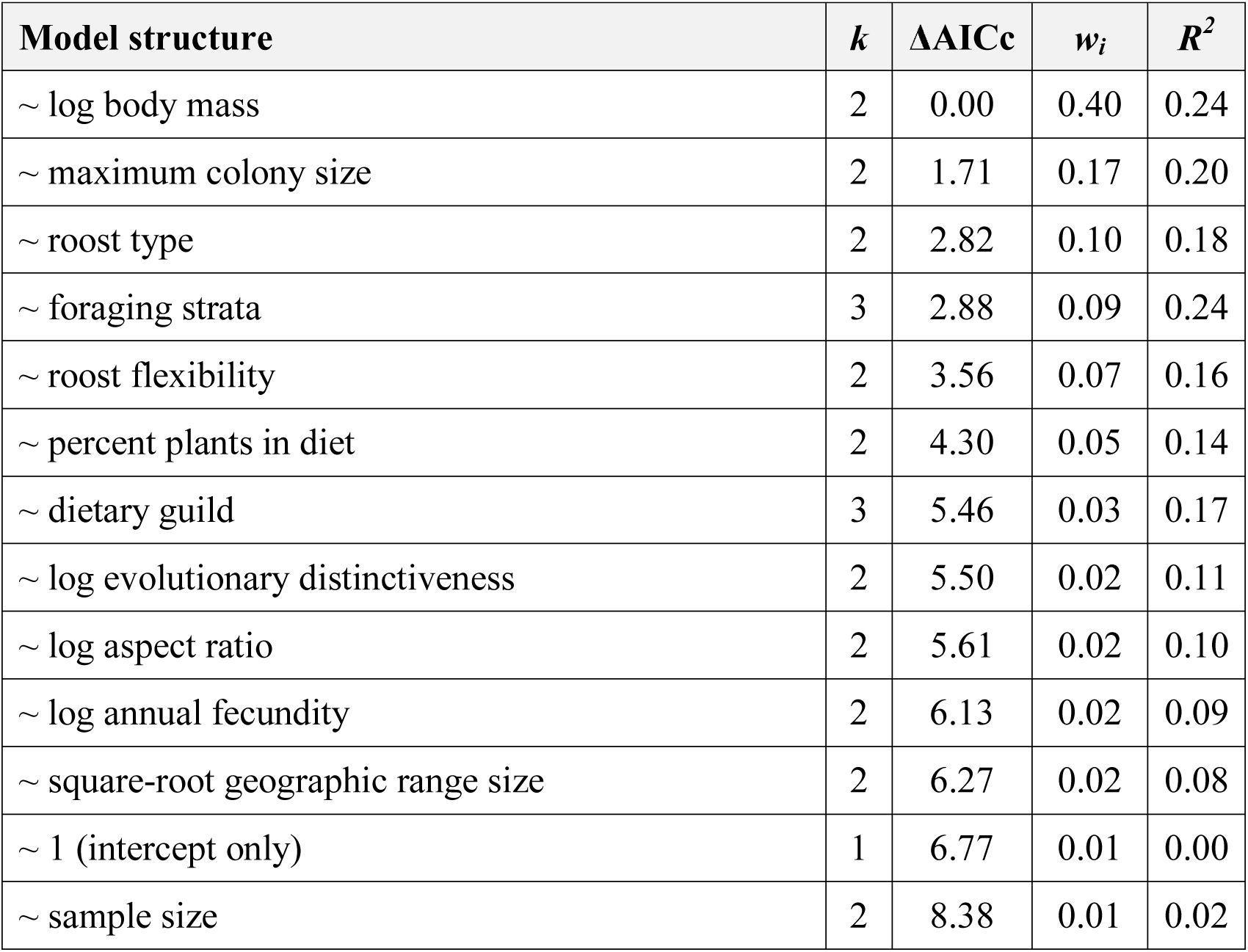
Competing weighted phylogenetic generalized least squares models predicting hemoplasma infection prevalence (logit-transformed) across the Belize bat community. Models are ranked by ΔAICc with the number of coefficients (*k*), Akaike weights (*w_i_*), and a likelihood ratio test pseudo-*R^2^*.

### Hemoplasma genotype diversity

Our phylogenetic analysis identified 29 *Mycoplasma* genotypes in the Belize bat community (Table 2), including three previously identified from vampire bats (VBG1–3; Volokhov et al., 2017). All genotypes demonstrated minor levels of sequence variability (99.5–100%; Table 2). Based on comparisons with sequences from GenBank, 21 of these genotypes represent novel hemoplasmas, five are closely related to non-hemoplasma mycoplasmas, and many are phylogenetically related to previously identified *Mycoplasma* spp. and hemoplasmas identified from other bat species (e.g., Di Cataldo et al., 2020; Millán et al., 2019, 2015), bat ticks (e.g., Hornok et al., 2019), primates including humans (e.g., Alcorn et al., 2020; Bonato, Figueiredo, Gonçalves, Machado, & André, 2015; Hattori et al., 2020), and rodents (e.g., Gonçalves et al., 2015; Goto, Yasuda, Hayashimoto, & Ebukuro, 2010; Vieira et al., 2009). A more detailed description of these 29 bacterial genotypes is provided in the Supporting Information (Fig. S3).

**Table 2.**
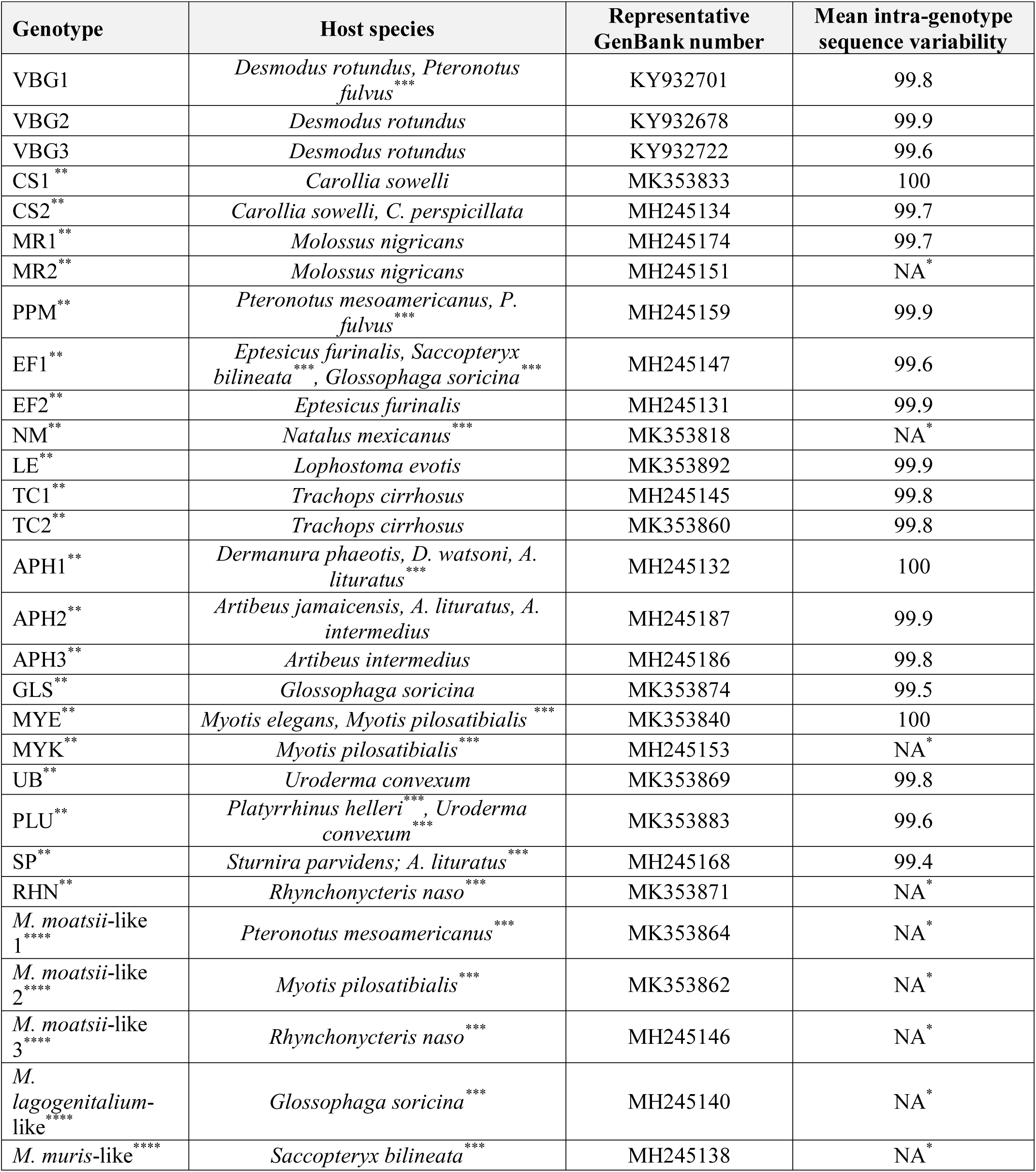
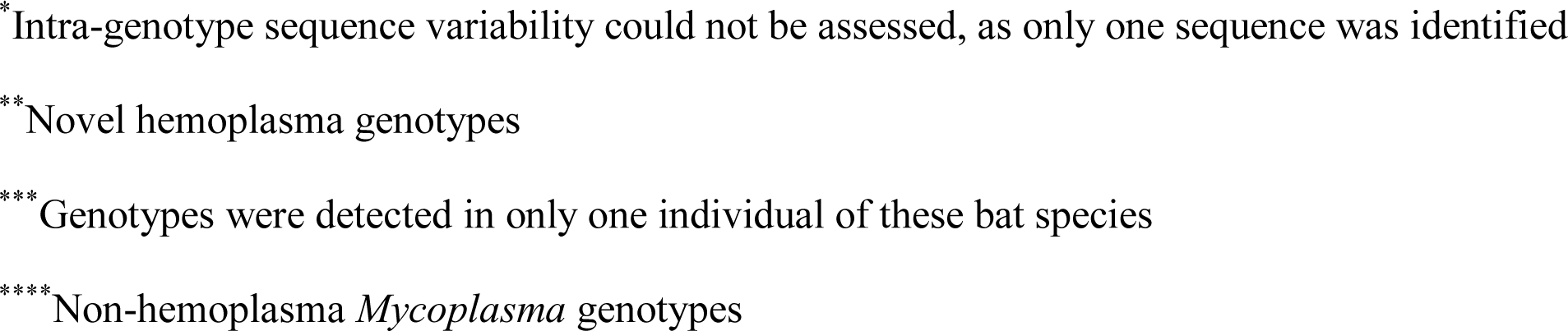
Hemoplasma genotypes identified from the Belize bat community. Genotypes are given with their bat host species, representative GenBank numbers, and intra-genotype variability.

| Genotype | Host species | Representative GenBank number | Mean intra-genotype sequence variability |
| --- | --- | --- | --- |
| VBG1 | *Desmodus rotundus*, *Pteronotus fulvus* *** | KY932701 | 99.8 |
| VBG2 | *Desmodus rotundus* | KY932678 | 99.9 |
| VBG3 | *Desmodus rotundus* | KY932722 | 99.6 |
| CS1 ** | *Carollia sowelli* | MK353833 | 100 |
| CS2 ** | *Carollia sowelli*, *C. perspicillata* | MH245134 | 99.7 |
| MR1 ** | *Molossus nigricans* | MH245174 | 99.7 |
| MR2 ** | *Molossus nigricans* | MH245151 | NA * |
| PPM ** | *Pteronotus mesoamericanus*, *P. fulvus* *** | MH245159 | 99.9 |
| EF1 ** | *Eptesicus furinalis*, *Saccopteryx bilineata* ***, *Glossophaga soricina* *** | MH245147 | 99.6 |
| EF2 ** | *Eptesicus furinalis* | MH245131 | 99.9 |
| NM ** | *Natalus mexicanus* *** | MK353818 | NA * |
| LE ** | *Lophostoma evotis* | MK353892 | 99.9 |
| TC1 ** | *Trachops cirrhosus* | MH245145 | 99.8 |
| TC2 ** | *Trachops cirrhosus* | MK353860 | 99.8 |
| APH1 ** | *Dermanura phaeotis*, *D. watsoni*, *A. lituratus* *** | MH245132 | 100 |
| APH2 ** | *Artibeus jamaicensis*, *A. lituratus*, *A. intermedius* | MH245187 | 99.9 |
| APH3 ** | *Artibeus intermedius* | MH245186 | 99.8 |
| GLS ** | *Glossophaga soricina* | MK353874 | 99.5 |
| MYE ** | *Myotis elegans*, *Myotis pilosatibialis* *** | MK353840 | 100 |
| MYK ** | *Myotis pilosatibialis* *** | MH245153 | NA * |
| UB ** | *Uroderma convexum* | MK353869 | 99.8 |
| PLU ** | *Platyrrhinus helleri* ***, *Uroderma convexum* *** | MK353883 | 99.6 |
| SP ** | *Sturnira parvidens*; *A. lituratus* *** | MH245168 | 99.4 |
| RHN ** | *Rhynchonycteris naso* *** | MK353871 | NA * |
| *M. moatsii*-like 1 **** | *Pteronotus mesoamericanus* *** | MK353864 | NA * |
| *M. moatsii*-like 2 **** | *Myotis pilosatibialis* *** | MK353862 | NA * |
| *M. moatsii*-like 3 **** | *Rhynchonycteris naso* *** | MH245146 | NA * |
| *M. lagogenitalium*-like **** | *Glossophaga soricina* *** | MH245140 | NA * |
| *M. muris*-like **** | *Saccopteryx bilineata* *** | MH245138 | NA * |

\*Intra-genotype sequence variability could not be assessed, as only one sequence was identified
\*\*Novel hemoplasma genotypes
\*\*\*Genotypes were detected in only one individual of these bat species
\*\*\*\*Non-hemoplasma *Mycoplasma* genotypes

After controlling for multiple comparisons, our 29 bacterial genotypes were associated with site (χ*^2^*=47.11, *p*<0.01) and year (χ*^2^*=40.40, *p*<0.01). Genotype composition was more diverse at LAR (Fig. S4), and KK hemoplasmas were dominated by vampire bat genotypes (VBG1–3). Genotype composition was more idiosyncratic by study year. However, these 29 bacterial genotypes were most strongly associated with bat species (χ*^2^*=3532, *p*<0.01; Fig. S4).

### Bat–hemoplasma evolutionary relationships

Although some hemoplasma genotypes were shared between bat species (i.e., VBG1, CS2, PPM, EF1, AH1–2, MYE, PLU, SP; *n*=9), most showed strong host specificity (*n*=20; Table 2). Our coevolutionary analysis (PACo) supported congruence between the bat and hemoplasma phylogenies (*m*^2^_XY_=36.21, *p*=0.02, *n=*1000; Fig. 4), suggesting that hemoplasma evolution has mostly tracked bat speciation. However, PACo also demonstrated that only 56% of the 41 unique bat–hemoplasma links displayed significant evidence of coevolution (Fig. S5), and these patterns were almost exclusively found within the Phyllostomidae (with the exception of *Saccopteryx bilineata* and its *Mycoplasma muris*–like bacterial genotype). The other 18 bat–hemoplasma links therefore displayed evidence of phylogenetic incongruence and thus likely host shifts.

**Figure 4.**
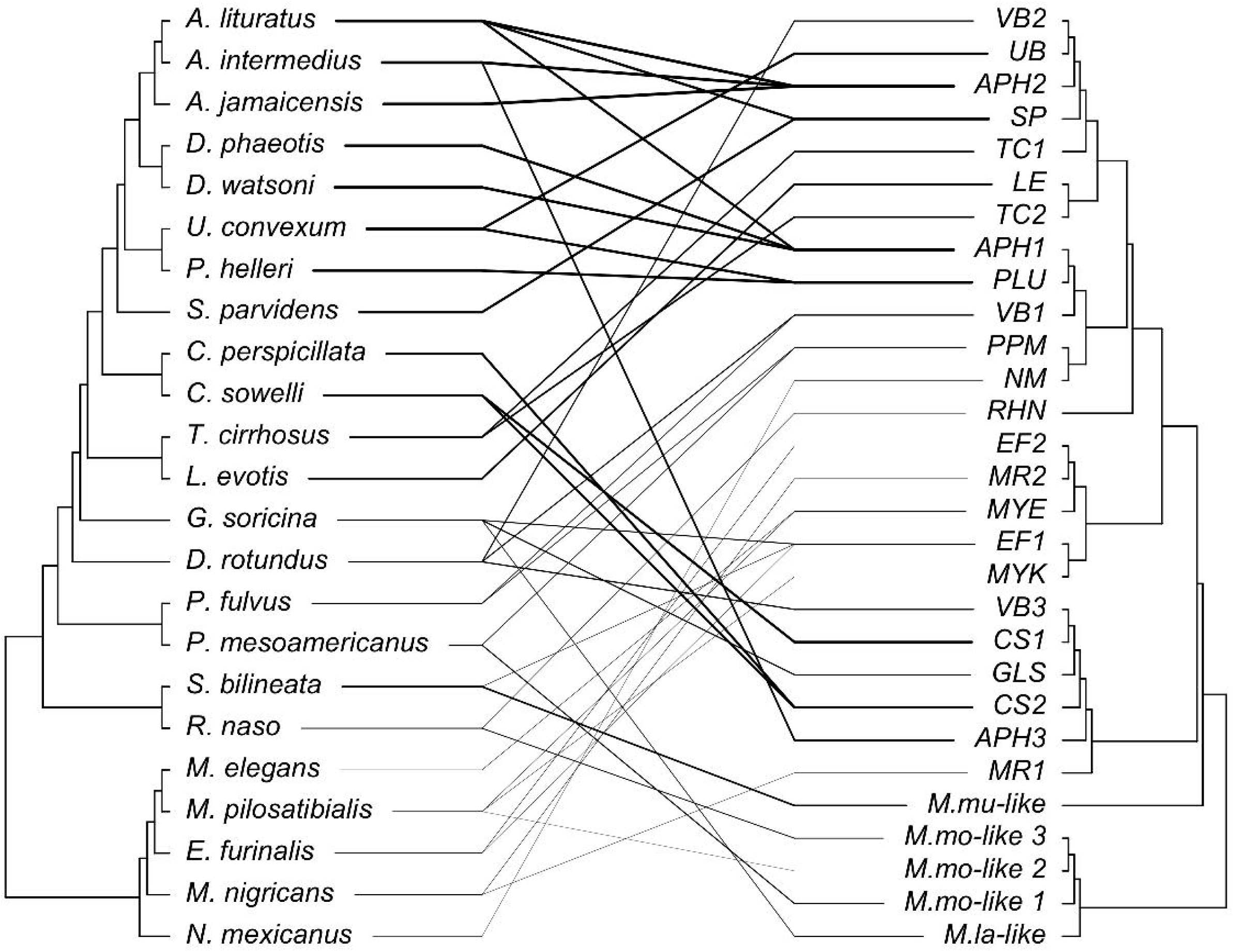
Evolutionary relationships between Belize bats and hemoplasma genotypes. The cophylogeny plot shows the bat phylogeny on the left and the hemoplasma genotype phylogeny on the right. We used the *treespace* package to collapse our complete hemoplasma phylogeny (Fig. S3) to only the 29 bacterial genotypes (Jombart, Kendall, Almagro Garcia, & Colijn, 2017). Lines display bat–hemoplasma associations and are shaded by the inverse of the squared residuals from PACo (i.e., dark lines show small residuals more indicative of coevolution).

### Hemoplasma genotype sharing networks

Within our bat–hemoplasma network, genotype sharing was restricted to five host communities, whereas six genotypes were each restricted to a single bat species (Fig. 5A). GLMs showed that both the frequency and presence of genotype sharing declined with phylogenetic distance between bat species (Poisson: *p*<0.001, *R^2^*=0.08; binomial: *p*<0.001, *R^2^*=0.51; Fig. 5B).

**Figure 5.**
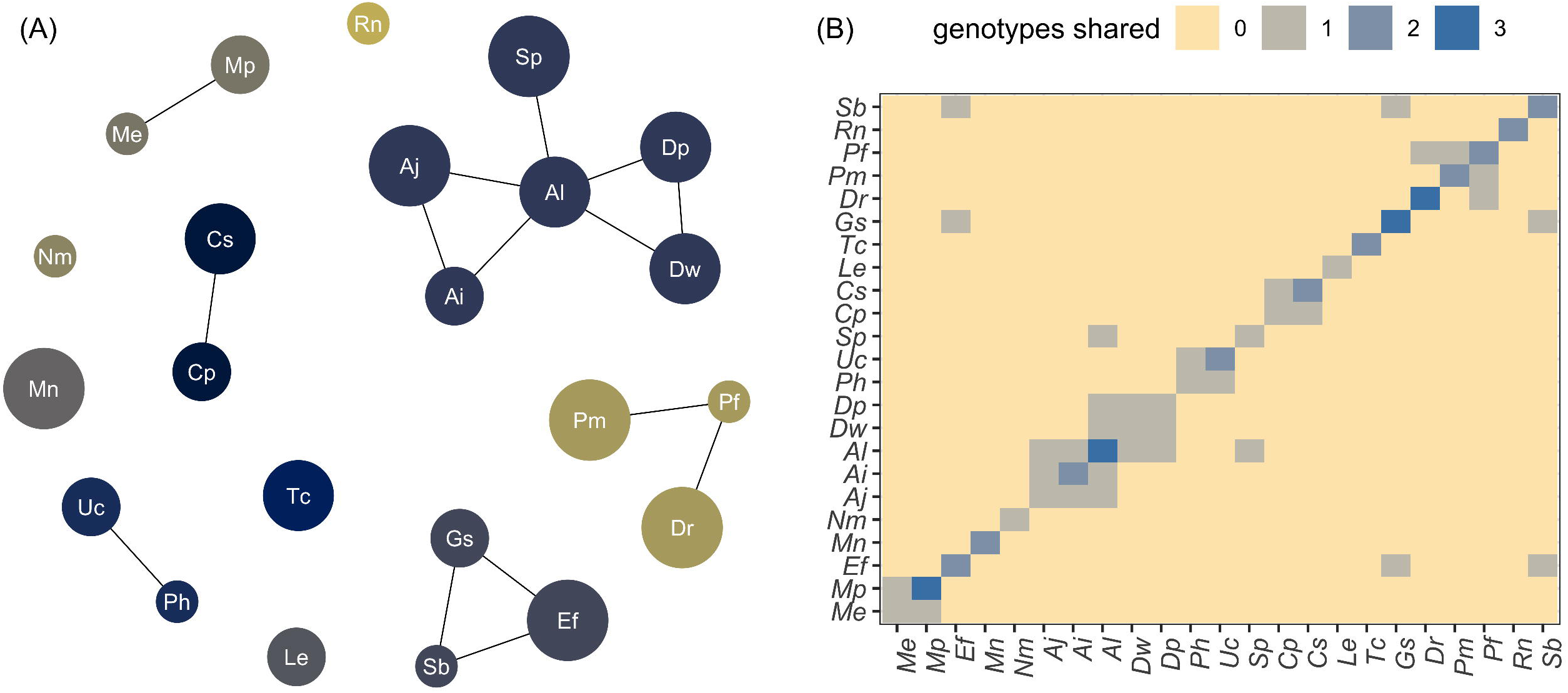
Patterns of hemoplasma genotype sharing across the Belize bat community. (A) Nodes in the genotype network represent bat species (abbreviated by Latin binomials), and edges represent a shared genotype. Nodes are colored by communities identified with the Louvain method and are scaled by the number of individuals per species. (B) The matrix shows pairwise hemoplasma genotype sharing, colored by the number of genotypes shared between bat species.

Bat species shared hemoplasma genotypes with zero to five other species (i.e., degree), and most hosts were not central to the network of genotype sharing (i.e., eigenvector centrality of zero). Six bat species had non-zero eigenvector centrality values that ranged from 37% to 100%, indicating that these hosts generally shared more hemoplasma genotypes with other highly connected hosts. Stratifying our hemoplasma genotype network across sites and years showed that centrality measures varied by space but not time (Fig. S7, Table S5). We observed no hemoplasma genotype sharing at KK, likely reflecting lower host diversity (Herrera et al., 2018).

Phylogenetic factorization identified similar bat clades with significantly different centrality compared to the paraphyletic remainder (Fig. 6A–B). For degree, the algorithm only identified *Artibeus lituratus* as being more central (x=5) than other bats (x=1.14). However, phylogenetic factorization identified three taxa in the subfamily Stenodermatinae that had significantly elevated eigenvector centrality: the genera *Artibeus* and *Dermanura* (x=0.67 compared to x=0.02 in all other bats), the species *Artibeus lituratus* (x=1 compared to x=0.12), and the species *Sturnira parvidens* (x=0.37 compared to x=0.15). Mirroring these results, phylogenetic signal was absent for degree (λ=0) but high for eigenvector centrality (λ=0.93).

**Figure 6.**
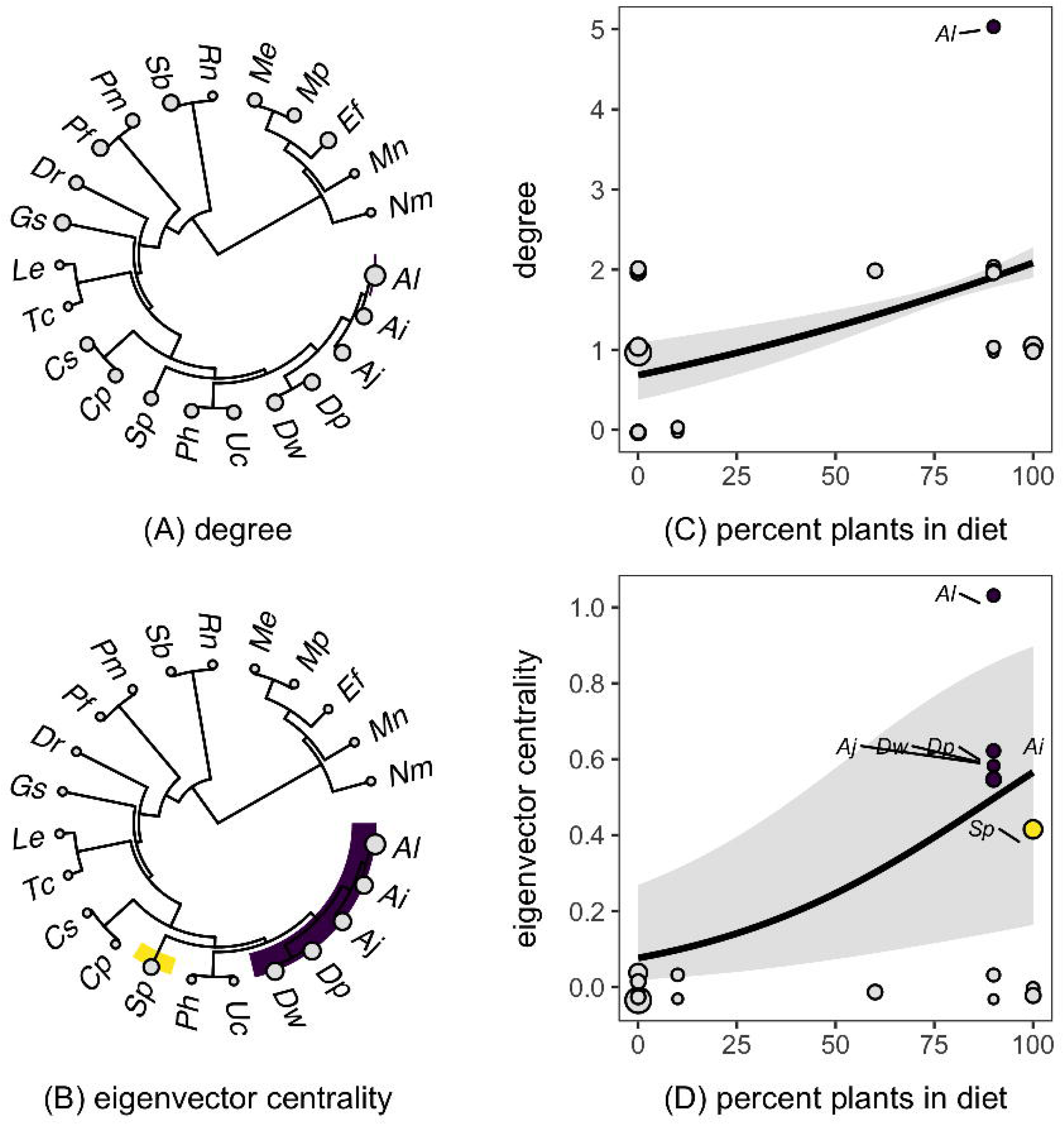
Phylogenetic patterns in hemoplasma genotype networks for Belize bat species (A) degree and (B) eigenvector centrality. Clades showing significantly different centrality metrics are highlighted, and points are scaled by observed values. Results from the top PGLS models predicting both centrality metrics as a function of bat species traits (C–D). Model fit and 95% confidence intervals are shown overlaid with data scaled by sample size; species from the clades identified through phylogenetic factorization are colored as in panels A–B.

Trait-based analyses showed that degree centrality was best predicted by diet (Table S6); bat species feeding more heavily on fruit and nectar shared more bacterial genotypes with other species (*β*=0.004, *p*<0.001, *R^2^*=0.20; Fig. 6C). Similarly, eigenvector centrality was best predicted by bat colony size and diet (Table S7); highly central species had small colonies (*β*_large_=–1.93, *p*=0.05, *R^2^*=0.13) and fed more on plants (*β*=0.03, *p*<0.01, *R^2^*=0.10; Fig. 6D).

As a final analysis, we assessed whether network centrality (i.e., a bat species’ role in hemoplasma genotype sharing) predicted contemporary infection prevalence (Fig. S8). However, we found no associations between species-level infection prevalence and centrality as measured by degree (*β* =–0.13, *R^2^*=0.03, *p=*0.42) or eigenvector centrality (*β*=–0.20, *R^2^*<0.01, *p*=0.79).

## Discussion

By examining the prevalence and distribution of a common bacterial pathogen (hemoplasmas) in a diverse bat community, we expanded analysis of the ecological and evolutionary predictors of bat infection and pathogen sharing beyond viruses. Across the bat community, hemoplasma infection risk was weakly higher for males but was better predicted by phylogeny, with large-bodied and large-colony bat species showing greater prevalence. Hemoplasmas showed high diversity and mostly strict host associations, with strong congruence between the bat and hemoplasma phylogenies. Although codivergence was supported by our analyses, we also observed hemoplasma genotype sharing and evidence of historical host shifts between closely related bats. Species most central to this hemoplasma sharing network displayed taxonomic clustering and were disproportionately frugivores and nectarivores. Yet these highly central bat species did not also have the highest hemoplasma prevalence, reinforcing mostly infrequent bacterial sharing between species. Our work reveals phylogenetic patterns in hemoplasma infection in a diverse bat community while contributing to broader efforts to understand the host specificity of bacterial pathogens and their cross-species transmission patterns in wildlife.

Whereas many bacterial pathogens, including hemoplasmas, are common in bats (Bai et al., 2011; Becker et al., 2018; Ikeda et al., 2017; Mascarelli et al., 2014; Millán et al., 2015; Volokhov et al., 2017), the factors that confer high infection probability are poorly understood. In the Belize bat community, the odds of hemoplasma infection tended to be higher in males, suggesting male-biased transmission as detected in feline and canine systems (Soto et al., 2017; Walker Vergara et al., 2016). Such patterns could stem from males mounting weaker immune responses than females (Kelly, Stoehr, Nunn, Smyth, & Prokop, 2018) or to male defense of multi-female roosts in many Neotropical bat species (Voigt, von Helversen, Michener, & Kunz, 2001). Direct transmission of hemoplasmas has been demonstrated in feline and rodent systems (Cohen et al., 2018; Museux et al., 2009) but only inferred in bats from metagenomic studies detecting these bacteria in saliva (Volokhov et al., 2017). We found weak support for the hypothesis that ectoparasites play a role in infection risks (Hornok et al., 2019; Willi, Boretti, Meli, et al., 2007). The weak male bias in infection could cast further doubt on vector-borne transmission, as females in some bat species have elevated ectoparasitism (Frank, Mendenhall, Judson, Daily, & Hadly, 2016); however, a secondary GLMM testing this hypothesis found generally weak support for a female bias in ectoparasitism in our system (Fig. S9). Future work assessing ectoparasite burdens could better elucidate the roles of vectors in hemoplasma risk.

Across Neotropical bats sampled in Belize, we found phylogeny to be a better predictor of hemoplasma risk than individual traits, site, or year. Phylogenetic factorization identified one clade, the Emballonuridae, with significantly lower prevalence than all other bats in the community. This moderate phylogenetic signal mirrors comparable effects of phylogeny for bat viruses (Guy, Thiagavel, Mideo, & Ratcliffe, 2019), similarly suggesting potential for innate differences in species susceptibility or pathogen exposure. Trait-based analyses revealed that this taxonomic pattern was driven by heavier and large-colony species having greater hemoplasma prevalence. Small-bodied species could have low prevalence due to small blood volumes and low bacterial titers (Volokhov et al., 2017). Alternatively, the positive, saturating relationship between body mass and bacterial prevalence could be driven by allometric patterns in competence (Downs et al., 2019), in contrast to weak or opposite relationships between mass and viral richness across bats (Guy et al., 2019; Han, Schmidt, et al., 2016). As larger-bodied bat species can also be more abundant in Neotropical habitat fragments (Herrera et al., 2018), these results suggest land conversion could increase the frequency of bat species most capable of maintaining hemoplasma infection. Similarly, positive relationships between colony size and prevalence could support density-dependent transmission of bacteria (McCallum et al., 2001), whereas mixed support has been found for bat viruses (Streicker et al., 2012; Webber, Fletcher, & Willis, 2017). Future work could test how community-wide infection patterns vary across broader habitat gradients and use multiple bacteria to assess the generality of these trends.

Approximately two-thirds of the Neotropical bat species sampled in Belize were infected by hemoplasmas, for which we observed high genetic diversity consistent with other studies of this pathogen in bats (Mascarelli et al., 2014; Millán et al., 2015; Volokhov et al., 2017). However, these bacterial genotypes were mostly novel and only weakly related to hemoplasmas described elsewhere in Latin America (Ikeda et al., 2017; Millán et al., 2019), with the exception of those previously identified from vampire bats (Volokhov et al., 2017). When considering the phylogenetic scale of genotypes, most hemoplasmas were host specific. Over half our hemoplasma communities consisted of a single bat–genotype association, matching the degree of host specificity observed more generally for *Mycoplasma* spp. (Citti & Blanchard, 2013; Pitcher & Nicholas, 2005). When we did detect genotype sharing between species, this occurred mostly between closely related hosts (e.g., PPM was detected in *Pteronotus mesoamericanus* and *P. fulvus*), indicating bat phylogenetic distance decreased the probability of bacterial transfer.

Analyses to characterize species centrality to the hemoplasma genotype sharing network showed that one species (*Artibeus lituratus*) and the subfamily Stenodermatinae played key roles. This clade, and especially the genera *Artibeus* and *Dermanura* (formerly all classified in *Artibeus*), was the only taxon with non-zero connectivity, and this pattern was reflected in fruit-and nectar-based diets and small colonies being the primary predictors of centrality. The strictly frugivorous Stenodermatinae represents a recent divergence in the Phyllostomidae (Botero-Castro et al., 2013), and high centrality of these species may indicate weaker phylogenetic barriers for bacterial transmission between hosts in this clade. Two other analyses reinforced infrequent and conserved hemoplasma sharing between species. First, phylogenetic patterns in prevalence were distinct from those in genotype sharing centrality (e.g., large-colony species had higher prevalence but lower connectivity), and prevalence accordingly did not predict centrality. Second, we found general congruence between bat and hemoplasma phylogenies. Although this shows codivergence is a strong evolutionary force, congruence can also stem from preferential jumps to closely related hosts (De Vienne et al., 2013). Though we cannot rule out that some host shifts may be artefacts of the limited resolution of both the phylogenies, our evolutionary analyses and genotype sharing results imply that hemoplasma host shifts are possible yet rare.

By sampling a diverse assemblage of bacterial genotypes in an ecologically and evolutionary rich host community, our work has broader implications for our understanding of disease emergence. Many bacterial pathogens are thought to be generalists and relatively unlikely to specialize in a novel host (Pedersen, Altizer, Poss, Cunningham, & Nunn, 2005; Woolhouse & Gowtage-Sequeria, 2005), in contrast to many viruses in which host shifts are more common owing to high mutation rates and short infectious periods (Geoghegan et al., 2017; Longdon, Brockhurst, Russell, Welch, & Jiggins, 2014). Recent theoretical work suggests host shift speciation may be less common for bacteria because of higher phenotypic plasticity (e.g., the ability to reside in diverse habitats) and a slower tempo of evolution (Bonneaud et al., 2019). Obligate reliance of *Mycoplasma* spp. on host cells and more chronic infections likely explains their propensity to specialize (Citti & Blanchard, 2013; Cohen et al., 2018). More broadly, however, using genetics to infer pathogen sharing, rather than coarser phylogenetic scales (e.g., species complexes or genera), is increasingly showing that many bacterial strains may be more host specific (Withenshaw, Devevey, Pedersen, & Fenton, 2016). The high specialism of bat hemoplasma genotypes thus underscores the importance of using finer phylogenetic scales in the study of infectious disease (Fountain-Jones et al., 2018; Graham, Storch, & Machac, 2018).

Comparative analyses of viruses have suggested that phylogenetically conserved pathogen jumps between species may be a broader generality in the study of disease emergence (Albery, Eskew, Ross, & Olival, 2019; Luis et al., 2015; Streicker et al., 2010). With few exceptions, our results on hemoplasma genotype sharing between Neotropical bat species are generally consistent with this pattern for a bacterial pathogen. Two cases in which hemoplasmas were shared between more distantly related species included the VBG1 genotype in *Desmodus rotundus* and *Pteronotus fulvus* (Phyllostomidae and Mormoopidae) and the EF1 genotype in *Glossophaga soricina* and *Saccopteryx bilineata* (Phyllostomidae and Emballonuridae). For the latter, both bat species co-roost in the LAR, which suggests an ecological context for pathogen exposure over current timescales. However, other genetic markers (e.g., *rpoB, rpoC, gyrB*) would be necessary to infer contemporary cross-species transmission (Kämpfer & Glaeser, 2012; Volokhov et al., 2012), as analysis of the 16s rRNA gene alone is insufficient for hemoplasma species identification (Volokhov et al., 2012). If hemoplasmas are more likely to specialize rather than expand their range into new and unrelated species, genotype sharing between unrelated bats could represent more transient spillovers (Bonneaud et al., 2019). As specialized pathogens could be more transmissible than generalists (Garamszegi, 2006), species with high infection prevalence of specialist genotypes could be prioritized for bacterial surveillance.

In conclusion, our analysis of a diverse community of bats and their pathogen genotypes identifies several key ecological and evolutionary factors structuring bacterial infection within and between species and provides a starting point for contrasts with such patterns for viruses. Similar to bat viruses, we found moderate phylogenetic signal in hemoplasma prevalence. However, these phylogenetic patterns in prevalence were decoupled from those describing bat species centrality in sharing hemoplasmas, such that genotype sharing was generally restricted by bat phylogeny. These findings imply codivergence of bats and their bacterial pathogens alongside rare and phylogenetically constrained host shifts. Future work more broadly characterizing the ecological and evolutionary determinants of bacterial infections in diverse host communities will improve our understanding of cross-species transmission beyond viruses and contribute to efforts to understand the epidemiological consequences of bacterial pathogens.

## Data accessibility

Individual-level data are available in Dryad (https://datadryad.org/stash/share/7qOGkejyIldAcD9FEZHNNVCc_4k0lmmiABy0EiRwZoo). Bat species trait data are available in Table S2. Hemoplasma sequences are available in GenBank (MH245119–MH245194 and MK353807–MK353892).

## Acknowledgements

For assistance with field logistics, bat sampling, and permits, we thank Mark Howells, Neil Duncan, James Herrera, Melissa Ingala, and staff of the Lamanai Field Research Center. We also thank the many colleagues who helped net bats during 2017–2018 bat research in Belize. We thank Seth Walk, Susan Broadway, and Jonathan Martinson for laboratory access and assistance. Lastly, we thank two anonymous reviewers for helpful suggestions to improve this manuscript.

## Statement of authorship

DJB, KAS, and AMB collected samples; NBS and MBF coordinated fieldwork; RKP, SA, and DGS supported laboratory analyses; DVV and VEC conducted molecular and phylogenetic analyses; and ADW and DGS assisted with statistical analyses. DJB analyzed data, produced figures, and wrote the manuscript. All authors provided critical review of the manuscript.

