## Supplemental Material for "Ecological and evolutionary drivers of hemoplasma infection and bacterial genotype sharing in a Neotropical bat community"

**S1. Bat sampling**

**S2. Comparative data**

**S3. Infection status and prevalence**

**S4. Hemoplasma genotypes**

**S5. Ectoparasitism**

### S1. Bat sampling

Table S1. Sample size per bat species included in analyses of hemoplasma infection, stratified by year and site (LAR and KK).

| | 2017 | | 2018 | |
|---|---|---|---|---|
| **Species** | **LAR** | **KK** | **LAR** | **KK** |
| *Artibeus intermedius* | 3 | 0 | 2 | 1 |
| *Artibeus jamaicensis* | 4 | 1 | 6 | 0 |
| *Artibeus lituratus* | 3 | 0 | 4 | 1 |
| *Bauerus dubiaquercus* | 4 | 0 | 2 | 0 |
| *Carollia perspicillata* | 3 | 2 | 2 | 1 |
| *Carollia sowelli* | 11 | 0 | 8 | 1 |
| *Chrotopterus auritus* | 0 | 3 | 1 | 0 |
| *Dermanura phaeotis* | 8 | 0 | 7 | 3 |
| *Dermanura watsoni* | 6 | 1 | 5 | 2 |
| *Desmodus rotundus* | 22 | 21 | 66 | 27 |
| *Eptesicus furinalis* | 8 | 1 | 10 | 0 |
| *Eumops nanus* | 1 | 0 | 0 | 0 |
| *Gardnerycteris keenani* | 0 | 0 | 2 | 0 |
| *Glossophaga soricina* | 11 | 0 | 7 | 0 |
| *Lasiurus ega* | 0 | 0 | 3 | 0 |
| *Lophostoma evotis* | 1 | 0 | 2 | 0 |
| *Micronycteris schmidtorum* | 1 | 0 | 0 | 0 |
| *Mimon cozumelae* | 1 | 0 | 0 | 5 |
| *Molossus nigricans* | 14 | 0 | 7 | 0 |
| *Mormoops megalophylla* | 0 | 0 | 1 | 1 |
| *Myotis elegans* | 0 | 0 | 2 | 0 |
| *Myotis pilosatibialis* | 1 | 0 | 3 | 0 |
| *Natalus mexicanus* | 0 | 0 | 1 | 0 |
| *Noctilio leporinus* | 0 | 0 | 1 | 0 |
| *Platyrrhinus helleri* | 0 | 0 | 1 | 1 |
| *Pteronotus fulvus* | 4 | 1 | 2 | 0 |
| *Pteronotus mesoamericanus* | 11 | 12 | 13 | 6 |
| *Rhogeesa aeneus* | 0 | 0 | 1 | 0 |
| *Rhynchonycteris naso* | 5 | 0 | 11 | 0 |
| *Saccopteryx bilineata* | 9 | 0 | 9 | 0 |
| *Sturnira parvidens* | 21 | 2 | 20 | 4 |
| *Trachops cirrhosus* | 0 | 3 | 4 | 1 |
| *Uroderma convexum* | 2 | 0 | 8 | 0 |

### S2. Comparative data

Figure S1. Pairwise phylogenetic distance between the 33 bat species included in our dataset.

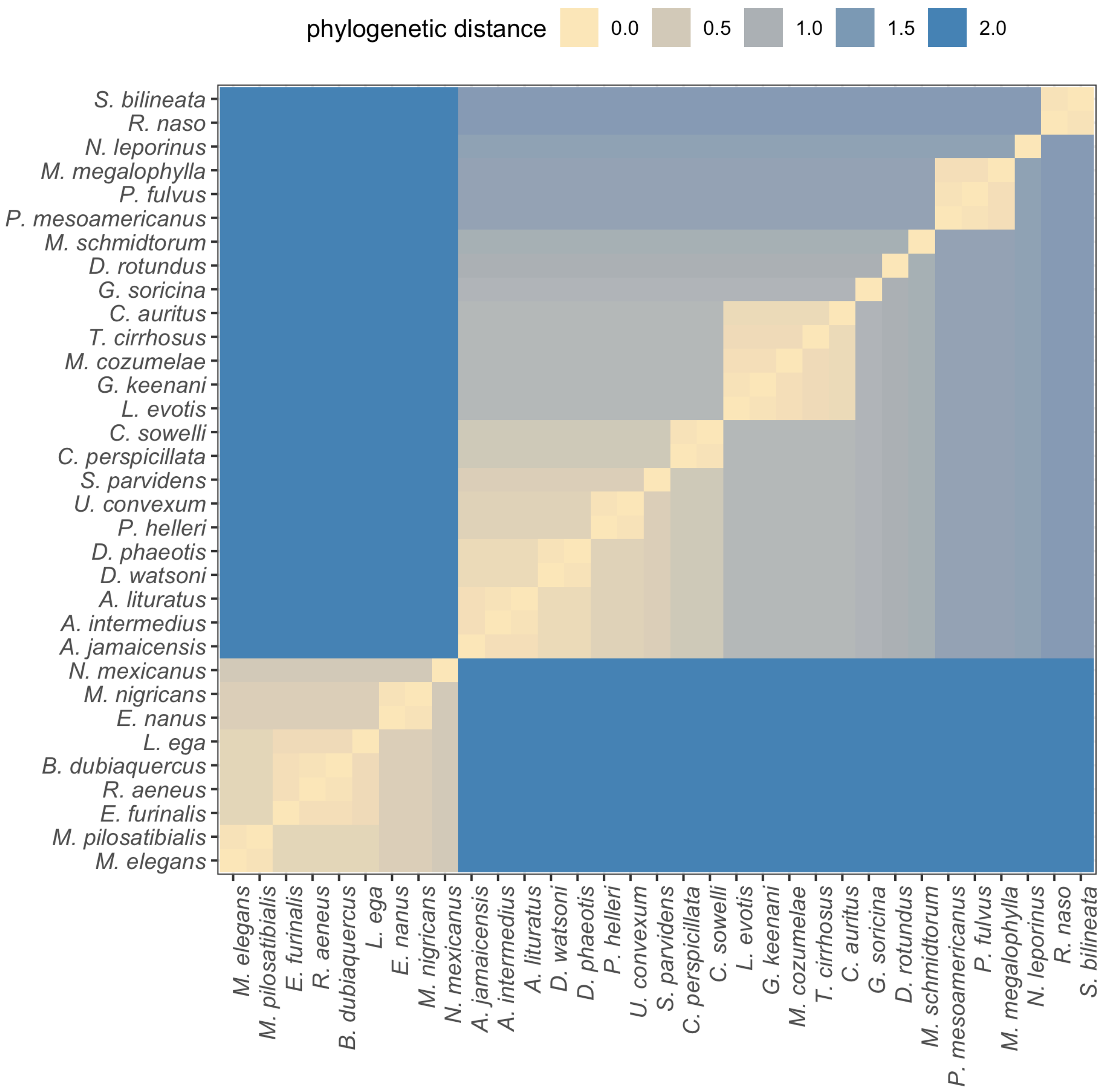

Figure S2. Map of geographic ranges for the 33 bat species sampled in Belize, stratified by whether hemoplasmas were detected. Transparent shapefiles from the International Union for Conservation of Nature are overlaid to highlight regions with dense distributional overlap.

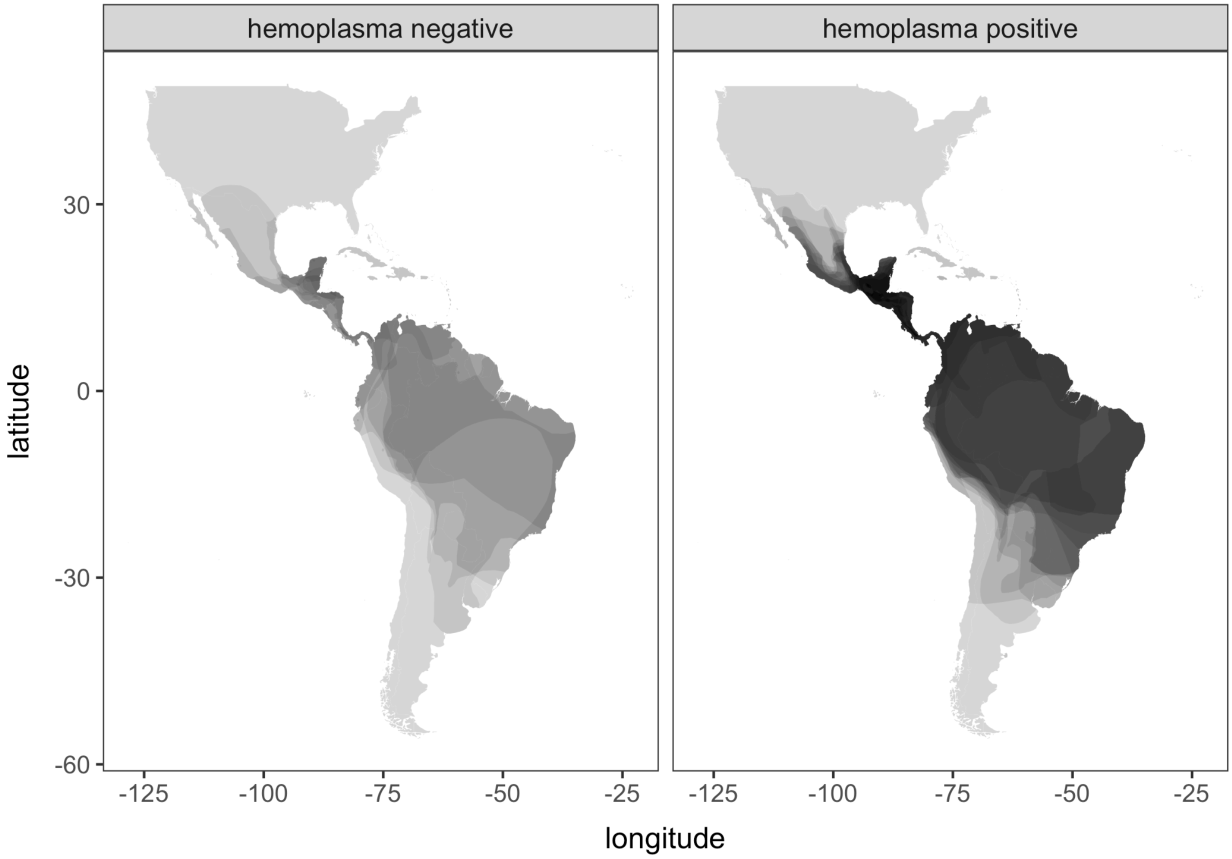

Table S2. Ecological and evolutionary trait data for the 33 Neotropical bat species included in phylogenetic comparative analyses.

| Species | | *N* | Mass | AF | Guild | % plant | Strata | AR | Roost | RF | Colony | $KM^2$ | ED |
|---|---|---|---|---|---|---|---|---|---|---|---|---|---|
| *Ai* | *Artibeus intermedius* | 6 | 60.5 | 1+ | plant | 90 | arboreal | 6.1 | closed | 1+ | small | 14235952 | 0.08 |
| *Aj* | *Artibeus jamaicensis* | 11 | 42.2 | 1+ | plant | 90 | arboreal | 6.4 | closed | 1+ | small | 1559896 | 0.12 |
| *Al* | *Artibeus lituratus* | 8 | 60.5 | 1+ | plant | 90 | arboreal | 6.1 | closed | 1+ | small | 14235952 | 0.08 |
| *Bd* | *Bauerus dubiaquercus* | 6 | 22.2 | 1+ | insect | 0 | aerial | 6.1 | closed | 1 | small | 265235 | 0.08 |
| *Cp* | *Carollia perspicillata* | 8 | 18 | 1+ | plant | 100 | arboreal | 6.1 | closed | 1+ | large | 13796364 | 0.20 |
| *Cs* | *Carollia sowelli* | 20 | 16.5 | 1+ | plant | 100 | arboreal | 5.5 | closed | 1+ | large | 790178 | 0.20 |
| *Ca* | *Chrotopterus auritus* | 4 | 77.7 | 1 | carnivore | 30 | arboreal | 5.5 | closed | 1+ | small | 13116793 | 0.30 |
| *Dp* | *Dermanura phaeotis* | 18 | 11.6 | 1+ | plant | 90 | arboreal | 6.3 | open | 1 | small | 3737367 | 0.11 |
| *Dw* | *Dermanura watsoni* | 14 | 11.2 | 1+ | plant | 90 | arboreal | 5.9 | open | 1 | small | 554677 | 0.11 |
| *Dr* | *Desmodus rotundus* | 139 | 31.2 | 1+ | carnivore | 0 | ground/aquatic | 6.7 | closed | 1+ | large | 17726518 | 0.56 |
| *Ef* | *Eptesicus furinalis* | 19 | 7.74 | 1+ | insect | 0 | aerial | 6.2 | closed | 1+ | large | 16292708 | 0.13 |
| *En* | *Eumops nanus* | 1 | 11.9 | 1 | insect | 0 | aerial | 8.8 | closed | 1 | small | 48804 | 0.23 |
| *Gk* | *Gardnerycteris keenani* | 2 | 13.8 | 1 | insect | 50 | arboreal | 8.3 | NA | 1 | small | 11136331 | 0.08 |
| *Gs* | *Glossophaga soricina* | 18 | 9.4 | 1+ | plant | 60 | arboreal | 6.4 | closed | 1+ | medium | 14858410 | 0.51 |
| *Lega* | *Lasiurus ega* | 3 | 13.2 | 1+ | insect | 0 | aerial | 7.9 | closed | 1+ | small | 13205183 | 0.19 |
| *Le* | *Lophostoma evotis* | 3 | 20.6 | 1 | insect | 10 | arboreal | 5.3 | closed | 1+ | small | 176593 | 0.08 |
| *Ms* | *Micronycteris schmidtorum* | 1 | 7.5 | 1 | insect | 20 | arboreal | 5.7 | closed | 1 | small | 6504802 | 0.61 |
| *Mc* | *Mimon cozumelae* | 6 | 18.6 | 1 | insect | 30 | arboreal | 8.3 | closed | 1+ | small | 803519 | 0.13 |
| *Mn* | *Molossus nigricans* | 21 | 25 | 1+ | insect | 0 | aerial | 11.1 | closed | 1 | large | 14800275 | 0.23 |
| *Mm* | *Mormoops megalophylla* | 2 | 15.8 | 1 | insect | 0 | aerial | 7.1 | closed | 1 | large | 3738686 | 0.38 |
| *Me* | *Myotis elegans* | 2 | 4.1 | 1 | insect | 0 | aerial | 6.4 | closed | 1 | large | 587280 | 0.16 |
| *Mp* | *Myotis pilosatibialis* | 4 | 5.3 | 1 | insect | 0 | aerial | 6.4 | closed | 1 | large | 2736294 | 0.16 |
| *Nm* | *Natalus mexicanus* | 1 | 5.5 | 1 | insect | 0 | aerial | 5.8 | closed | 1 | large | 2433276 | 0.63 |
| *Nl* | *Noctilio leporinus* | 1 | 58.3 | 1 | carnivore | 0 | ground/aquatic | 9 | closed | 1+ | large | 14625648 | 0.76 |
| *Ph* | *Platyrrhinus helleri* | 2 | 13.5 | 1+ | plant | 90 | arboreal | 6.4 | closed | 1+ | small | 10016822 | 0.14 |
| *Pf* | *Pteronotus fulvus* | 7 | 9.21 | 1 | insect | 0 | aerial | 8.3 | NA | 1 | large | 3471860 | 0.20 |
| *Pm* | *Pteronotus mesoamericanus* | 42 | 19.7 | 1 | insect | 0 | aerial | 6.7 | closed | 1+ | large | 539713 | 0.20 |
| *Ra* | *Rhogeessa aeneus* | 1 | 4.2 | 1+ | insect | 0 | aerial | 6.1 | closed | 1 | large | 144232 | 0.08 |
| *Rn* | *Rhynchonycteris naso* | 16 | 3.8 | 1+ | insect | 0 | aerial | 6.5 | open | 1 | medium | 11508828 | 0.45 |
| *Sb* | *Saccopteryx bilineata* | 18 | 7.8 | 1 | insect | 0 | aerial | 6.1 | open | 1 | medium | 12632095 | 0.45 |
| *Sp* | *Sturnira parvidens* | 47 | 19.1 | 1+ | plant | 100 | arboreal | 6.5 | open | 1 | small | 4859514 | 0.30 |
| *Tc* | *Trachops cirrhosus* | 8 | 37.67 | 1 | carnivore | 10 | ground/aquatic | 6.3 | closed | 1+ | small | 12642872 | 0.20 |
| *Uc* | *Uroderma convexum* | 10 | 16.69 | 1+ | plant | 90 | arboreal | 6.3 | NA | 1 | medium | 12796601 | 0.14 |

[*]*N*: Number of individuals per bat species screened for hemoplasma infection; AF: annual fecundity; AR: aspect ratio; RF: roost flexibility; ED: evolutionary distinctiveness. Data for *Pf*, *Sp*, *Mn*, *Mp*, and *Uc* were taken from the bat species from which they were originally classified (*Pteronotus davyi*, *Sturnira lilium*, *Molossus rufus, Myotis keaysi*, and *Uroderma bilobatum*).

**S3. Infection status and prevalence**

Table S3. Hemoplasma infection prevalence (i.e., the proportion of sequence-confirmed, hemoplasma-positive bats) for each species, stratified by site and year in northern Belize.

| Species | Site | Year | *N* | + | % |
|---|---|---|---|---|---|
| *Artibeus intermedius* | KK | 2018 | 1 | 1 | 1 |
| *Artibeus intermedius* | LAR | 2017 | 3 | 2 | 0.67 |
| *Artibeus intermedius* | LAR | 2018 | 2 | 1 | 0.5 |
| *Artibeus jamaicensis* | KK | 2017 | 1 | 1 | 1 |
| *Artibeus jamaicensis* | LAR | 2017 | 4 | 4 | 1 |
| *Artibeus jamaicensis* | LAR | 2018 | 6 | 4 | 0.67 |
| *Artibeus lituratus* | KK | 2018 | 1 | 0 | 0 |
| *Artibeus lituratus* | LAR | 2017 | 3 | 2 | 0.67 |
| *Artibeus lituratus* | LAR | 2018 | 4 | 3 | 0.75 |
| *Bauerus dubiaquercus* | LAR | 2017 | 4 | 0 | 0 |
| *Bauerus dubiaquercus* | LAR | 2018 | 2 | 0 | 0 |
| *Carollia perspicillata* | KK | 2017 | 2 | 1 | 0.5 |
| *Carollia perspicillata* | KK | 2018 | 1 | 0 | 0 |
| *Carollia perspicillata* | LAR | 2017 | 3 | 1 | 0.33 |
| *Carollia perspicillata* | LAR | 2018 | 2 | 1 | 0.5 |
| *Carollia sowelli* | KK | 2018 | 1 | 1 | 1 |
| *Carollia sowelli* | LAR | 2017 | 11 | 2 | 0.18 |
| *Carollia sowelli* | LAR | 2018 | 8 | 4 | 0.5 |
| *Chrotopterus auritus* | KK | 2017 | 3 | 0 | 0 |
| *Chrotopterus auritus* | LAR | 2018 | 1 | 0 | 0 |
| *Dermanura phaeotis* | KK | 2018 | 3 | 0 | 0 |
| *Dermanura phaeotis* | LAR | 2017 | 9 | 4 | 0.44 |
| *Dermanura phaeotis* | LAR | 2018 | 7 | 4 | 0.57 |
| *Dermanura watsoni* | KK | 2017 | 1 | 1 | 1 |
| *Dermanura watsoni* | KK | 2018 | 2 | 1 | 0.5 |
| *Dermanura watsoni* | LAR | 2017 | 6 | 3 | 0.5 |
| *Dermanura watsoni* | LAR | 2018 | 5 | 0 | 0 |
| *Desmodus rotundus* | KK | 2017 | 21 | 20 | 0.95 |
| *Desmodus rotundus* | KK | 2018 | 29 | 14 | 0.48 |
| *Desmodus rotundus* | LAR | 2017 | 22 | 13 | 0.59 |
| *Desmodus rotundus* | LAR | 2018 | 68 | 38 | 0.56 |
| *Eptesicus furinalis* | KK | 2017 | 1 | 1 | 1 |
| *Eptesicus furinalis* | LAR | 2017 | 8 | 6 | 0.75 |
| *Eptesicus furinalis* | LAR | 2018 | 10 | 8 | 0.8 |
| *Eumops nanus* | LAR | 2017 | 1 | 0 | 0 |
| *Gardnerycteris keenani* | LAR | 2018 | 2 | 0 | 0 |
| *Glossophaga soricina* | LAR | 2017 | 11 | 2 | 0.18 |
| *Glossophaga soricina* | LAR | 2018 | 7 | 2 | 0.29 |

| | | | | | |
|---|---|---|---|---|---|
| *Lasiurus ega* | LAR | 2018 | 3 | 0 | 0 |
| *Lophostoma evotis* | LAR | 2017 | 1 | 1 | 1 |
| *Lophostoma evotis* | LAR | 2018 | 2 | 2 | 1 |
| *Micronycteris schmidtorum* | LAR | 2017 | 1 | 0 | 0 |
| *Mimon cozumelae* | KK | 2018 | 5 | 0 | 0 |
| *Mimon cozumelae* | LAR | 2017 | 1 | 0 | 0 |
| *Molossus nigricans* | LAR | 2017 | 14 | 13 | 0.93 |
| *Molossus nigricans* | LAR | 2018 | 7 | 3 | 0.43 |
| *Mormoops megalophylla* | KK | 2018 | 1 | 0 | 0 |
| *Mormoops megalophylla* | LAR | 2018 | 1 | 0 | 0 |
| *Myotis elegans* | LAR | 2018 | 2 | 2 | 1 |
| *Myotis pilosatibialis* | LAR | 2017 | 1 | 1 | 1 |
| *Myotis pilosatibialis* | LAR | 2018 | 3 | 2 | 0.67 |
| *Natalus mexicanus* | LAR | 2018 | 1 | 1 | 1 |
| *Noctilio leporhinus* | LAR | 2018 | 1 | 0 | 0 |
| *Platyrrhinus helleri* | KK | 2018 | 1 | 0 | 0 |
| *Platyrrhinus helleri* | LAR | 2018 | 1 | 1 | 1 |
| *Pteronotus fulvus* | KK | 2017 | 1 | 0 | 0 |
| *Pteronotus fulvus* | LAR | 2017 | 4 | 1 | 0.25 |
| *Pteronotus fulvus* | LAR | 2018 | 2 | 1 | 0.5 |
| *Pteronotus mesoamericanus* | KK | 2017 | 12 | 7 | 0.58 |
| *Pteronotus mesoamericanus* | KK | 2018 | 6 | 5 | 0.83 |
| *Pteronotus mesoamericanus* | LAR | 2017 | 11 | 8 | 0.73 |
| *Pteronotus mesoamericanus* | LAR | 2018 | 13 | 9 | 0.69 |
| *Rhogeesa aeneus* | LAR | 2018 | 1 | 0 | 0 |
| *Rhynchonycteris naso* | LAR | 2017 | 5 | 1 | 0.2 |
| *Rhynchonycteris naso* | LAR | 2018 | 11 | 1 | 0.09 |
| *Saccopteryx bilineata* | LAR | 2017 | 9 | 1 | 0.11 |
| *Saccopteryx bilineata* | LAR | 2018 | 9 | 1 | 0.11 |
| *Sturnira parvidens* | KK | 2017 | 2 | 1 | 0.5 |
| *Sturnira parvidens* | KK | 2018 | 4 | 2 | 0.5 |
| *Sturnira parvidens* | LAR | 2017 | 23 | 11 | 0.48 |
| *Sturnira parvidens* | LAR | 2018 | 20 | 13 | 0.65 |
| *Trachops cirrhosus* | KK | 2017 | 3 | 2 | 0.67 |
| *Trachops cirrhosus* | KK | 2018 | 1 | 1 | 1 |
| *Trachops cirrhosus* | LAR | 2018 | 4 | 4 | 1 |
| *Uroderma convexum* | LAR | 2017 | 2 | 1 | 0.5 |
| *Uroderma convexum* | LAR | 2018 | 8 | 3 | 0.38 |

Table S4. Competing phylogenetic GLMMs predicting hemoplasma infection status across the Belize bat community ($n$=323 after removing missing values). Models are ranked by ΔLOOIC with their LOOIC SE, Akaike weights ($w_i$) and Bayesian $R^2$ estimates.

<table>
<tr><th>Model structure</th><th>LOOIC</th><th>SE</th><th>ΔLOOIC</th><th>$w_i$</th><th>$R^2_m$</th><th>$R^2_c$</th></tr>
<tr><td>~ sex + ectoparasite + reproduction + site * year + (1|species) + (1|phylogeny)</td><td>404.56</td><td>16.1</td><td>0.00</td><td>0.29</td><td>0.07</td><td>0.23</td></tr>
<tr><td>~ sex + ectoparasite + reproduction + year + (1|species) + (1|phylogeny)</td><td>404.72</td><td>15.2</td><td>0.16</td><td>0.27</td><td>0.05</td><td>0.22</td></tr>
<tr><td>~ sex + ectoparasite + reproduction + site + year + (1|species) + (1|phylogeny)</td><td>405.77</td><td>15.3</td><td>1.21</td><td>0.16</td><td>0.05</td><td>0.22</td></tr>
<tr><td>~ sex * reproduction + ectoparasite + year + (1|species) + (1|phylogeny)</td><td>406.57</td><td>15.3</td><td>2.00</td><td>0.11</td><td>0.05</td><td>0.22</td></tr>
<tr><td>~ sex * reproduction + ectoparasite + site * year + (1|species) + (1|phylogeny)</td><td>406.72</td><td>16.2</td><td>2.16</td><td>0.10</td><td>0.07</td><td>0.24</td></tr>
<tr><td>~ sex * reproduction + ectoparasite + year + (1|species) + (1|phylogeny)</td><td>407.48</td><td>15.6</td><td>2.92</td><td>0.07</td><td>0.06</td><td>0.22</td></tr>
</table>

**S4. Hemoplasma genotypes**

We here provide details of how each identified genotype (Table 2) is phylogenetically related to one another and other sequences identified in GenBank (updated March 2020, see Fig. S3).

The MR1 genotype, identified from *Molossus nigricans* (formerly classified as *M. rufus* [1]), showed 92.5% and 94.3% similarity to *Candidatus* Mycoplasma turicensis and *Mycoplasma coccoides*, as well as 98.3% similarity to hemoplasmas previously detected in *M. molossus* in Brazil (KY356748) [2]. The PPM genotype, which had 98.6% similarity to VBG1 [3], was identified from *Pteronotus mesoamericanus* and in one sample from *P. fulvus* (formerly classified as *P. davyi* [4,5]; MK353825). The 16S rRNA sequence from another *P. fulvus* (MH245133) was identical to VBG1. The NM genotype, identified in one *Natalus mexicanus*, showed 99.3% and 98.3% similarity to PPM and VBG1, respectively; however, the species-specific pattern of mutations allowed us to discriminate NM from these two genotypes. The RHN genotype, identified in a single *Rhynchonycteris naso*, demonstrated a unique sequence with 93.3-94.4% similarity to *Candidatus* Mycoplasma haemohominis [6], *Candidatus* Mycoplasma turicensis, and *M. haemomuris*; 94.8% similarity to *Candidatus* Mycoplasma haemomacaque; and 95.6-96.1% similarity to the PPM and SP genotypes (see below).

Two distinct hemoplasma genotypes (CS1 and CS2) were identified in *Carollia* spp. CS1, only identified in *C. sowelli*, was 98.0% similar to VBG3, while CS2 (identified in both *C. sowelli* and *C. perspicillata*) was 97.7% and 97.4% similar to VBG3 and CS1, respectively. The unique GLS genotype identified in a single *Glossophaga soricina* demonstrated 98.1% similarity to VBG3 and 96.8% and 97.4% similarity to the CS1 and CS2 genotypes, respectively.

The EF1 genotype comprised hemoplasma sequences that were mainly identified in *Eptesicus furinalis* alongside single positive samples from *Saccopteryx bilineata* (MK353889) and *Glossophaga soricina* (MH245128); this genotype was 96.9% similar to *Candidatus* M. haemohominis as well as 96.7-98.2% similar to hemoplasma genotypes previously detected in *Miniopterus schreibersii* and *Myotis capaccinii* from Spain [7] and to a hemoplasma genotype previously detected in bat ticks (*Ixodes simplex*) from Hungary [8]. The MYK genotype, identified in one *Myotis pilosatibialis* (previously classified as *M. keaysi* [9]), was 97.3% similar to the EF1 genotype , 96.2% similar to *Candidatus* M. haemohominis, and 96.5-98.0% similar to the above-mentioned genotypes previously detected in bats from Spain [7] and bat ticks from Hungary [8]. Both the EF2 genotype (identified in *E. furinalis*) and the MR2 genotype (identified in *Molossus nigricans*) were 98.9% similar to one another; however, species-specific mutation patterns again facilitated discriminating these genotypes. Both genotypes were 97.7% similar to the MYE genotype from *Myotis elegans* and *M. pilosatibialis*, and all three of these genotypes (MYE, EF2, and MR2) showed 97.6-98.3% similarity to the hemoplasma genotype previously identified in little brown bats (*M. lucifugus*) from the eastern and northeastern United States [10] and to hemoplasmas detected in *M. chiloensis* from Chile (MK295627, MK295629, and MK295630) [11]. The EF2 and MR2 genotypes were also 93.5-94.5% similar to *Candidatus* M. haemomacaque, *M. haemomuris, Candidatus* M. turicensis, and *Candidatus* M. haemohominis.

The APH1 genotype was identified in *Dermanura phaeotis, D. watsoni*, and in one *A. lituratus*; this genotype showed 99.0% similarity to the PLU genotype detected in one *Platyrrhinus helleri* and in one *Uroderma convexum* (formerly classified as *U. bilobatum* [12,13]). These genotypes were further discriminated based on species-specific pattern of their mutations. The APH1 and PLU genotypes were 96.3-96.6% similar to the APH2 genotype from *Artibeus jamaicensis, A. lituratus*, and *A. intermedius*. The APH2 genotype showed 98.3%

similarity to VBG2 [3] and the UB genotype. The APH3 genotype was found in *A. intermedius* and was 96.2-96.7% similar to the CS1 and CS2 genotypes.

Two genotypes (TC1 and TC2) identified in *Trachops cirrhosus* were 96.7% similar to each other, and both were 96.6% similar to the LE genotype identified in *Lophostoma evotis*. The TC1, TC2, and LE genotypes were 97.3%, 96.3%, and 96.6% similar to the PPM and NM genotypes. The SP genotype, which comprised slightly divergent (99.4% intra-genotype sequence similarity) but closely related hemoplasma species, was identified in *Sturnira parvidens* (previously classified as *S. lilium* [14,15]) and in one *A. lituratus*. The SP genotype was most closely related to the UB genotype (99.1% similar) and was 98.4% similar to the APH2 genotype and VBG2. However, due to some diversity of SP sequences and their similarity to UB, APH2, and VBG2, the SP genotype did not form a distinct branch with significant bootstrap support on the phylogeny (Fig. S3). Thus, sequences within the SP genotype may comprise at least three closely related hemoplasma species or geographic clones (or isolates) of the same species (e.g., see the hypothetical groups A, B, and C on Fig. S3). However, definitive conclusions on the hemoplasma species repertoire within the SP genotype cannot be made given only analysis of the partial 16S rRNA gene; the nucleotide and phylogenetic analysis of additional housekeeping genes (e.g., *rpoB, gyrB*) [16,17] or the employment of next-generation sequencing methods [18–20] to screen for the full spectrum of blood-borne bacteria in bat samples will be required for future studies of these hemoplasma genotypes. All genotypes demonstrated minor levels of intra-genotype sequence variability (99.4–100%; see Table 2).

Lastly, three non-hemotropic *Mycoplasma* spp. genotypes were identified from five different bat species in Belize. Three *M. moatsii*–like sequences (numbered as 1–3) from *Pteronotus mesoamericanus, Myotis pilosatibialis*, and *Rhynchonycteris naso* showed 95.2%,

98.1%, and 96.9% similarity, respectively, to the 16S rRNA gene of *M. moatsii* type strain MK405 (NR_025186), a non-hemotropic *Mycoplasma sp.* isolated from grivet monkeys (*Cercopithecus aethiops*) [21]. We previously reported the *M. moatsii*–like genotype in *Desmodus rotundus* (KY932724) from Belize sampled in 2014 and 2015 [3]. However, these three novel *M. moatsii*–like sequences were not identical to each other or to the *M. moatsii*–like genotype previously found in *D. rotundus* (numbered as 4)*;* thus, the *M. moatsii*–like genotype 1 showed 97.0%, 96.1%, and 96.9% similarity to the *M. moatsii*–like genotypes 2-4, the *M. moatsii*–like genotype 2 showed 97.0%, 97.9%, and 98.2% similarity to the *M. moatsii*–like genotypes 1, 3-4, the *M. moatsii*–like genotype 3 showed 96.1%, 97.9%, and 96.3% similarity to the *M. moatsii*–like genotypes 1, 2, 4. This suggests these sequences are derived from closely related but different non-hemotropic *Mycoplasma* spp. in bats. The *M. lagogenitalium*-like genotype from *G. soricina* showed 96.5% and 97.6% similarity to the 16S rRNA genes of *M. lagogenitalium* type strain 12MS (NR_025185), a non-hemotropic *Mycoplasma sp.* described in Afghan pikas (*Ochotona rufescens*) [22] and *Mycoplasma* sp. EDS-4 isolated from house musk shrew (*Suncus murinus*) [23]. The *M. muris*–like genotype identified from *Saccopteryx bilineata* demonstrated 98.1% similarity to the 16S rRNA genes of *M. muris* type strain RIII-4 (NR_044664), a non-hemotropic *Mycoplasma* sp. isolated from mice [24].

*References*


1. Loureiro LO, Engstrom MD, Lim BK. 2020 Single nucleotide polymorphisms (SNPs) provide unprecedented resolution of species boundaries, phylogenetic relationships, and genetic diversity in the mastiff bats (Molossus). *Mol. Phylogenet. Evol.* **143**, 106690.

2. Ikeda P *et al.* 2017 Evidence and molecular characterization of <span class="italic">Bartonella</span> spp. and hemoplasmas in neotropical bats in Brazil. *Epidemiol. Amp Infect.* , 1–15. (doi:10.1017/S0950268817000966)

3. Volokhov DV, Becker DJ, Bergner LM, Camus MS, Orton RJ, Chizhikov VE, Altizer SM, Streicker DG. 2017 Novel hemotropic mycoplasmas are widespread and genetically diverse in vampire bats. *Epidemiol. Infect.*

12. Mantilla-Meluk H. 2014 *Defining species and species boundaries in Uroderma (Chiroptera: Phyllostomidae) with a description of a new species*. Museum of Texas Tech University.

Figure S3. Phylogenetic relationships based on sequence data for the partial 16S rRNA gene (approx. 864-878 bp) among the hemoplasma genotypes detected in Belize bat species with other hemotropic and non-hemotropic *Mycoplasma* spp. Accession numbers are shown alongside bat species names. The tree was constructed with the minimum evolution method in MEGA X. Bootstrap values were evaluated from 1000 replications; those < 50% are not included on the tree. *Streptococcus equi* is used as an outgroup. The bar indicates 0.05 substitutions per site.

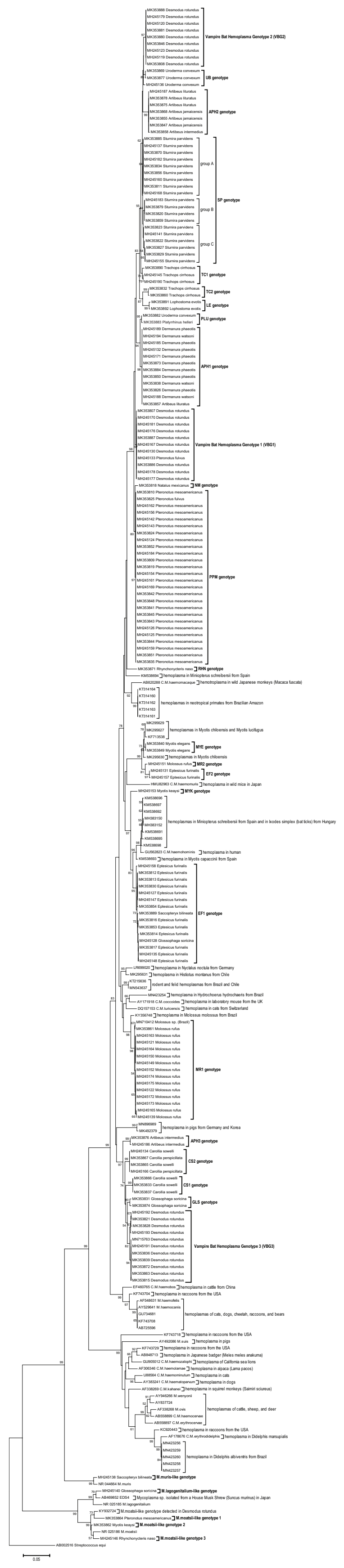

Figure S4. Relative abundance of the 29 hemoplasma genotypes detected in the Belize Neotropical bat community in northern Belize according to site, year, and bat species.

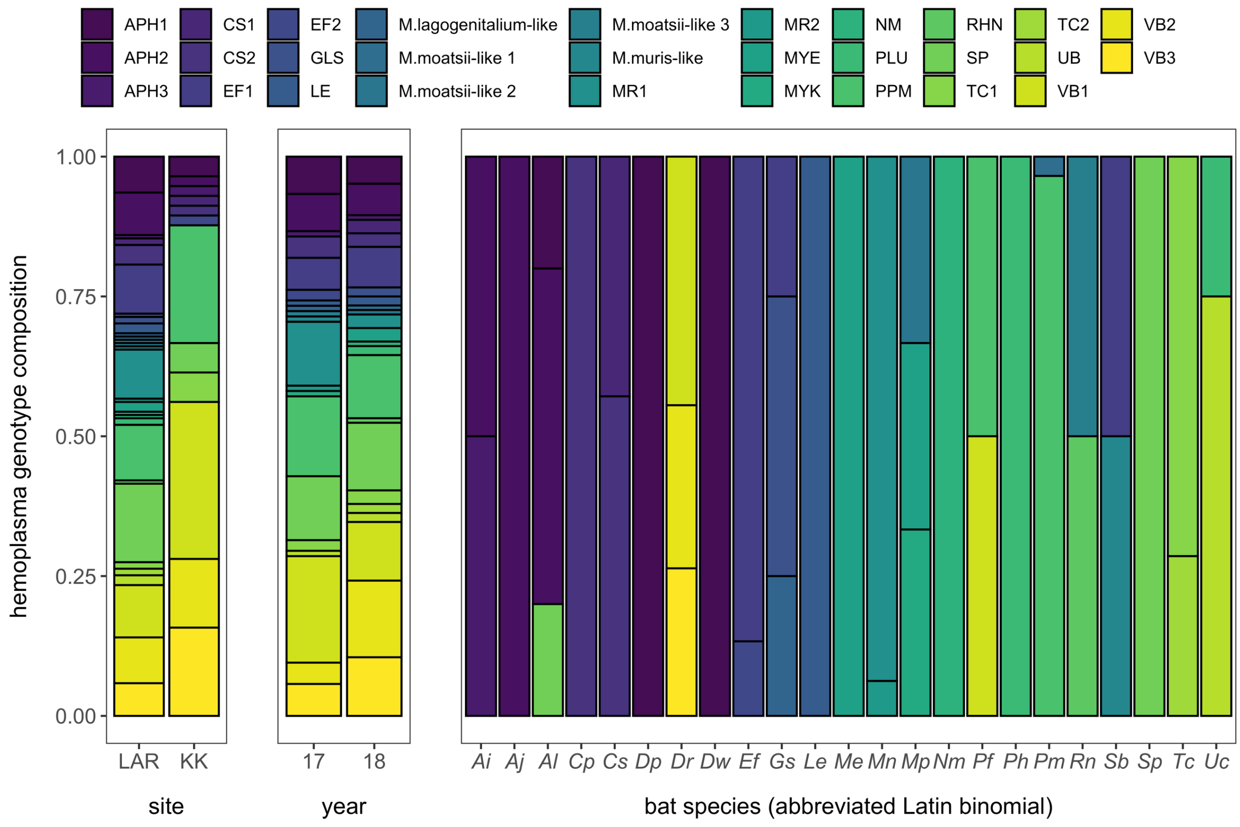

Figure S5. Contribution of unique bat–genotype links (squared residuals) to cophylogenetic signal ($m^2_{XY}$) from PACo. Links are considered supportive of coevolution if the upper 95% confidence interval falls below the mean squared residuals (shown in the dashed line).

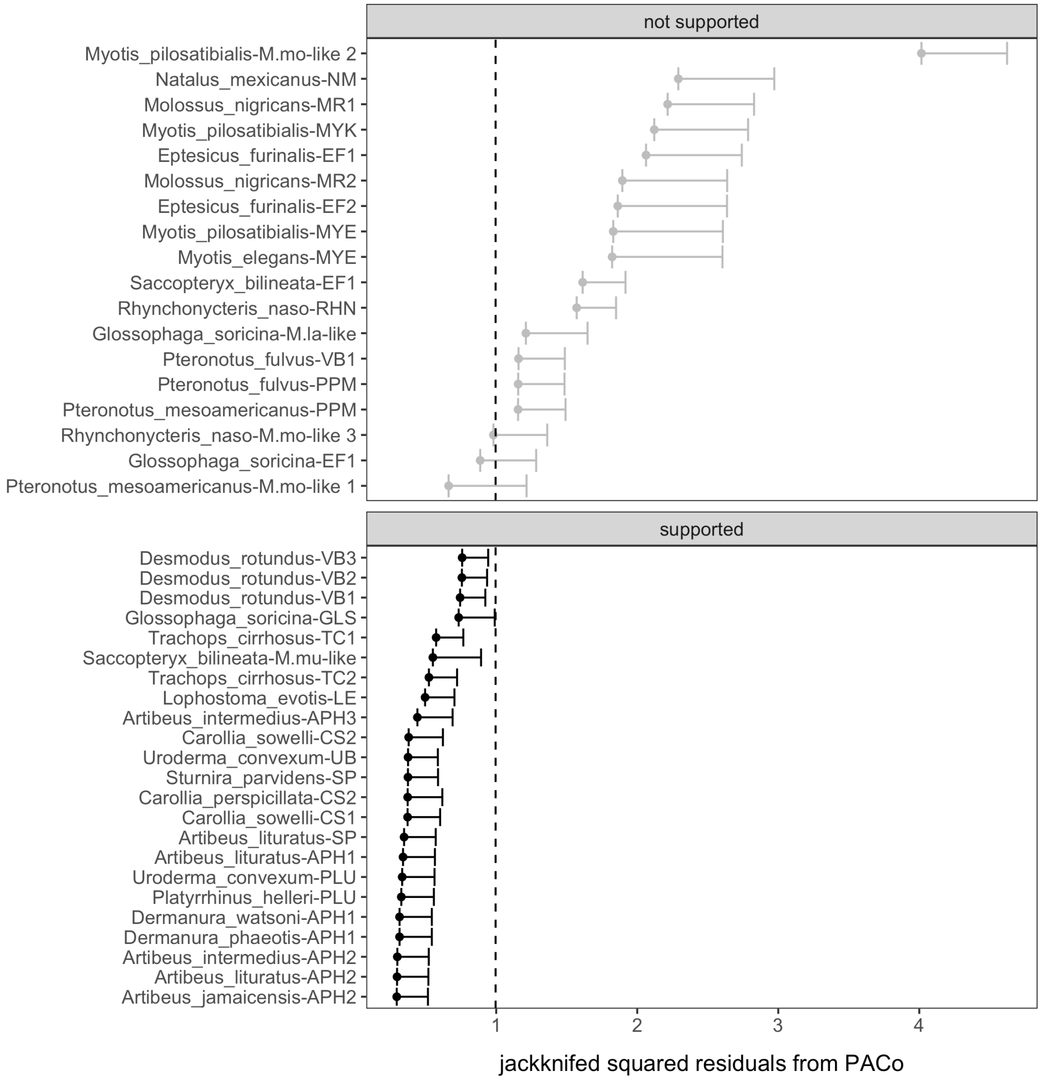

Figure S6. Metrics of network centrality (degree, eigenvector centrality) for sharing of hemoplasma genotypes between bat species in Belize.

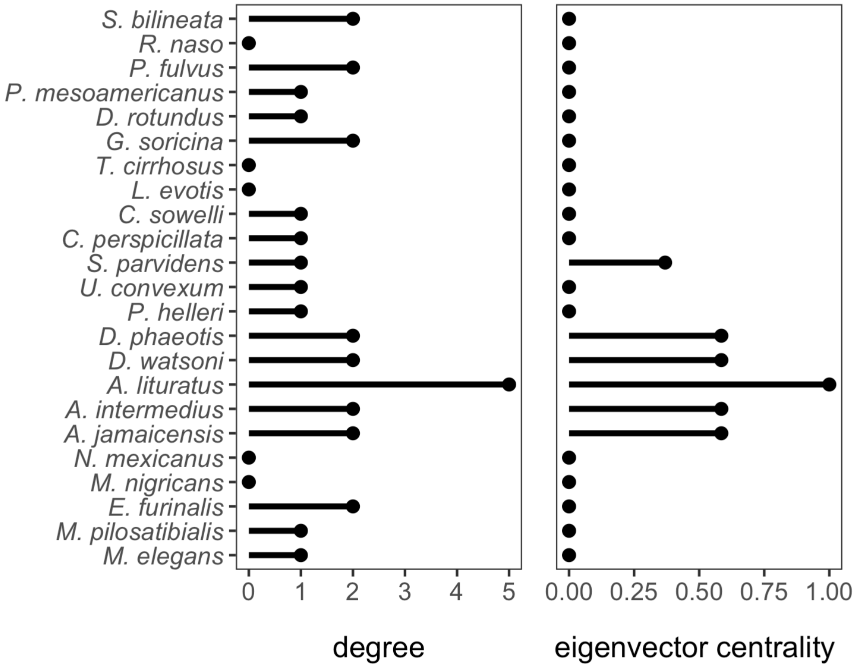

Figure S7. Networks of hemoplasma genotype sharing across the Belize bat community, stratified by site and year. Nodes represent bat species (abbreviated by their Latin binomial), and edges represent whether two nodes share any hemoplasma genotypes. Nodes with no edges represent bat species that do not share any hemoplasma genotypes with other bat species.

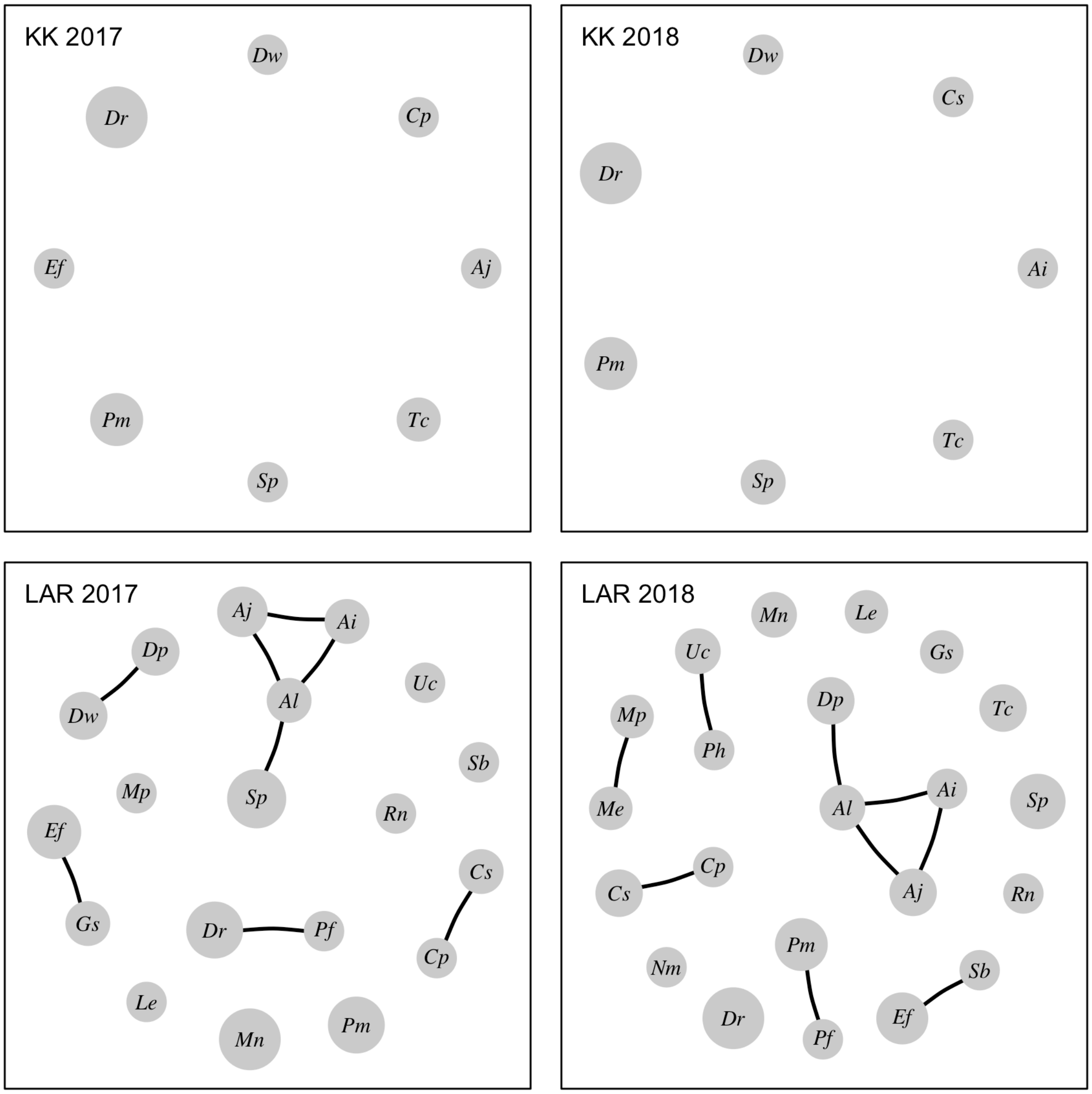

Table S5. Results of GLMs applied to each hemoplasma genotype sharing network centrality measure as a function of site, year, and their interaction term.

| **Variable** | $\chi^2$ | ***p*** |
|---|---|---|
| *Degree* | | |
| Site | 21.19 | <0.001 |
| Year | 0.007 | 0.93 |
| Site * year | 0 | 1 |
| *Eigenvector centrality* | | |
| Site | 112.33 | <0.001 |
| Year | 0.046 | 0.83 |
| Site * year | 0.017 | 0.90 |

Table S6. Competing PGLS models predicting degree (from the hemoplasma genotype network) across the Belize bat community. Models are ranked by ΔAICc with the number of coefficients (*k*), Akaike weights ($w_i$), and a likelihood ratio test pseudo-$R^2$.

| **Model structure** | ***k*** | **ΔAICc** | $w_i$ | $R^2$ |
|---|---|---|---|---|
| Square-root degree ~ % plants | 2 | 0 | 0.25 | 0.21 |
| Square-root degree ~ log aspect ratio | 2 | 0.82 | 0.16 | 0.18 |
| Square-root degree ~ dietary guild | 3 | 1.66 | 0.11 | 0.25 |
| Square-root degree ~ log annual fecundity | 2 | 2.02 | 0.09 | 0.14 |
| Square-root degree ~ log evolutionary distinctiveness | 2 | 2.12 | 0.08 | 0.13 |
| Square-root degree ~ 1 | 1 | 2.77 | 0.06 | 0 |
| Square-root degree ~ roost flexibility | 2 | 3.35 | 0.05 | 0.09 |
| Square-root degree ~ foraging strata | 3 | 3.51 | 0.04 | 0.19 |
| Square-root degree ~ colony size | 2 | 3.59 | 0.04 | 0.08 |
| Square-root degree ~ log mass | 2 | 3.76 | 0.04 | 0.07 |
| Square-root degree ~ square-root geographic range | 2 | 4.09 | 0.03 | 0.06 |
| Square-root degree ~ roost type | 2 | 4.1 | 0.03 | 0.06 |
| Square-root degree ~ sample size | 2 | 5.42 | 0.02 | 0 |

Table S7. Competing PGLS models predicting eigenvector centrality (from the hemoplasma genotype network) across the Belize bat community. Models are ranked by ΔAICc with the number of coefficients (*k*), Akaike weights ($w_i$), and a likelihood ratio test pseudo-$R^2$.

| **Model structure** | ***k*** | **ΔAICc** | $w_i$ | $R^2$ |
|---|---|---|---|---|
| Logit eigenvector centrality ~ colony size | 2 | 0 | 0.24 | 0.13 |
| Logit eigenvector centrality ~ 1 | 1 | 0.61 | 0.18 | 0 |
| Logit eigenvector centrality ~ % plants | 2 | 0.87 | 0.16 | 0.1 |
| Logit eigenvector centrality ~ log evolutionary distinctiveness | 2 | 2.22 | 0.08 | 0.04 |
| Logit eigenvector centrality ~ roost type | 2 | 3.04 | 0.05 | 0.01 |
| Logit eigenvector centrality ~ log mass | 2 | 3.21 | 0.05 | 0 |
| Logit eigenvector centrality ~ sample size | 2 | 3.27 | 0.05 | 0 |
| Logit eigenvector centrality ~ roost flexibility | 2 | 3.28 | 0.05 | 0 |
| Logit eigenvector centrality ~ log annual fecundity | 2 | 3.65 | 0.04 | 0 |
| Logit eigenvector centrality ~ square-root geographic range | 2 | 3.8 | 0.04 | 0 |
| Logit eigenvector centrality ~ log aspect ratio | 2 | 3.82 | 0.04 | 0 |
| Logit eigenvector centrality ~ dietary guild | 3 | 4.34 | 0.03 | 0.08 |
| Logit eigenvector centrality ~ foraging strata | 3 | 4.87 | 0.02 | 0.06 |

Figure S8. Associations between hemoplasma genotype sharing network centrality metrics and hemoplasma infection prevalence (logit-transformed as the response variable). PGLS model fit and 95% confidence intervals are shown overlaid with bat species data scaled by sample size.

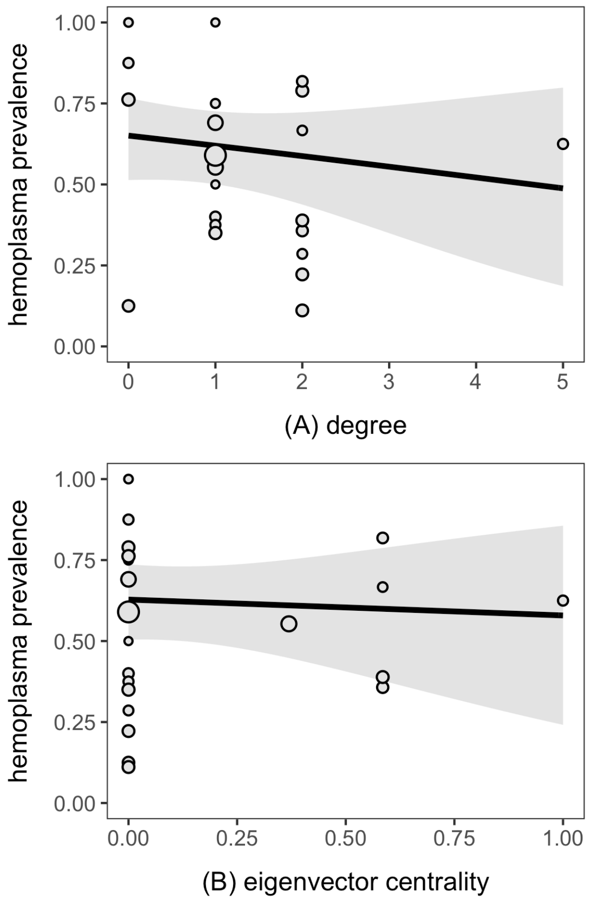

**S5. Ectoparasitism**

Figure S9. Predictors of individual ectoparasite presence. Odds ratios and 95% HDIs from the phylogenetic GLMMs, including the univariate sex effect and that adjusted for other covariates. Reference levels include males, bats sampled at LAR, reproductive bats, and bats from 2017.

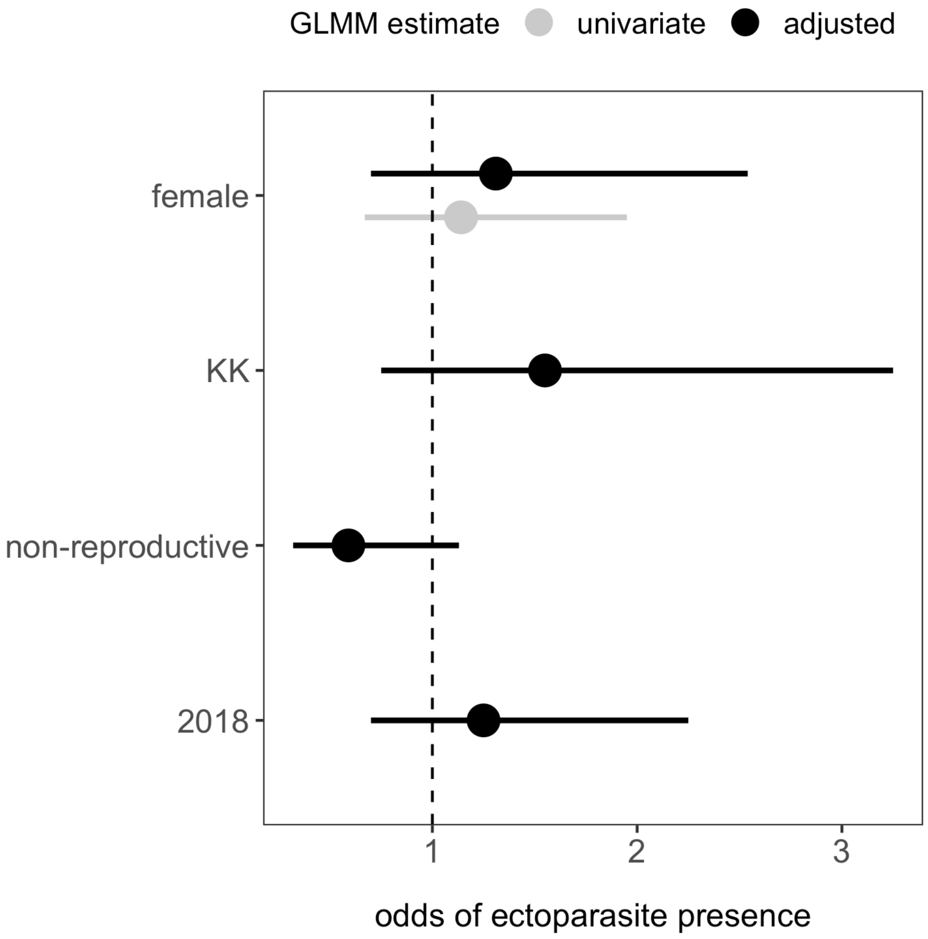
